# ipmr: Flexibly implement Integral Projection Models in R

**DOI:** 10.1101/2021.04.20.440590

**Authors:** Sam C. Levin, Dylan Z. Childs, Aldo Compagnoni, Sanne Evers, Tiffany M. Knight, Roberto Salguero-Gómez

## Abstract

1. Integral projection models (IPMs) are an important tool for studying the dynamics of populations structured by one or more continuous traits (*e.g.* size, height, color). Researchers use IPMs to investigate questions ranging from linking drivers to plant population dynamics, planning conservation and management strategies, and quantifying selective pressures in natural populations. The popularity of stage-structured population models has been supported by *R* scripts and packages (*e.g.* `IPMpack, popbio, popdemo, lefko3`) aimed at ecologists, which have introduced a broad repertoire of functionality and outputs. However, pressing ecological, evolutionary, and conservation biology topics require developing more complex IPMs, and considerably more expertise to implement them. Here, we introduce `ipmr`, a flexible *R* package for building, analyzing, and interpreting IPMs.
2. The `ipmr` framework relies on the mathematical notation of the models to express them in code format. Additionally, this package decouples the model parameterization step from the model implementation step. The latter point substantially increases `ipmr`’s flexibility to model complex life cycles and demographic processes.
3. `ipmr` can handle a wide variety of models, including density dependence, discretely and continuously varying stochastic environments, and multiple continuous and/or discrete traits. `ipmr` can accommodate models with individuals cross-classified by age and size. Furthermore, the package provides methods for demographic analyses (*e.g.* asymptotic and stochastic growth rates) and visualization (*e.g.* kernel plotting).
4. `ipmr` is a flexible *R* package for integral projection models. The package substantially reduces the amount of time required to implement general IPMs. We also provide extensive documentation with six vignettes and help files, accessible from an *R* session and online.

## Introduction

Integral projection models (IPMs) are an important and widely used tool for ecologists studying structured population dynamics in discrete time. Since the paper introducing IPMs over two decades ago (Easterling et al., 2000), at least 255 peer-reviewed publications on at least 250 plant species and 60 animal species have used IPMs (Table S1, Fig. S1). These models have addressed questions ranging from invasive species population dynamics (*e.g.* Crandall & Knight, 2017), effect of climate drivers on population persistence (*e.g.* Compagnoni et al., 2020), evolutionary stable strategies (*e.g.* Childs et al., 2004), and rare/endangered species conservation (*e.g.* Ferrer-Cervantes et al., 2012).

The IPM was introduced as alternative to matrix population models, which model populations structured by discrete traits (Caswell, 2001). Some of the advantages of using an IPM include (i) the ability to model populations structured by continuously distributed traits, (ii) the ability to flexibly incorporate discrete and continuous traits in the same model (*e.g.* seeds in a seedbank and a height structured plant population (Crandall & Knight, 2017), or number of females, males, and age-1 recruits for fish species (Erickson et al., 2017)), (iii) efficient parameterization of demographic processes with familiar regression methods (Coulson, 2012), and (iv) the numerical discretization of continuous kernels (see below) means that the tools available for matrix population models are usually also applicable for IPMs. Furthermore, researchers have developed methods to incorporate spatial dynamics (Jongejans et al., 2011), environmental stochasticity (Rees & Ellner, 2009), and density/frequency dependence into IPMs (Adler et al., 2010, Ellner et al., 2016). These developments were accompanied by the creation of software tools and guides to assist with IPM parameterization, implementation, and analysis. These tools range from *R* scripts with detailed annotations (Coulson, 2012, Merow et al., 2014, Ellner et al., 2016) to *R* packages (Metcalf et al., 2013, Shefferson et al., 2020).

Despite the array of resources available to researchers, implementing an IPM is still not a straightforward exercise. For example, an IPM that simulates a population for 100 time steps requires the user to either write or adapt from published guides multiple functions (*e.g.* to summarize demographic functions into the proper format), implement the numerical approximations of the model’s integrals, ensure that individuals are not accidentally sent beyond the integration bounds (“unintentional eviction”, *sensu* Williams et al., 2012), and track how the population state changes over the course of a simulation. Stochastic IPMs present further implementation challenges. In addition to the aforementioned elements, users must generate the sequence of environments that the population experiences. There are multiple ways of simulating environmental stochasticity, each with their own strengths and weaknesses (Metcalf et al. 2015).

`ipmr` manages these key details while providing the user flexibility in their models. `ipmr` uses the `rlang` package for metaprogramming (Henry & Wickham, 2019), which enables `ipmr` to provide a miniature domain specific language for implementing IPMs. `ipmr` aims to mimic the mathematical syntax that describes IPMs as closely as possible (Fig. 1, Tables 1 and 2). This *R* package can handle models with individuals classified by a mixture of any number of continuously and discretely distributed traits. Furthermore, `ipmr` introduces specific classes and methods to deal with both discrete and continuous stochastic environments, density-independent and -dependent models, as well as age-structured populations. `ipmr` decouples the parameterization (*i.e.* regression model fitting) and implementation steps (*i.e.* converting the regression parameters into a full IPM), and does not attempt to help users with the parameterization task. This provides greater flexibility in modeling trait-demography relationships, and enables users to specify IPMs of any functional form that they desire.

**Figure 1:**
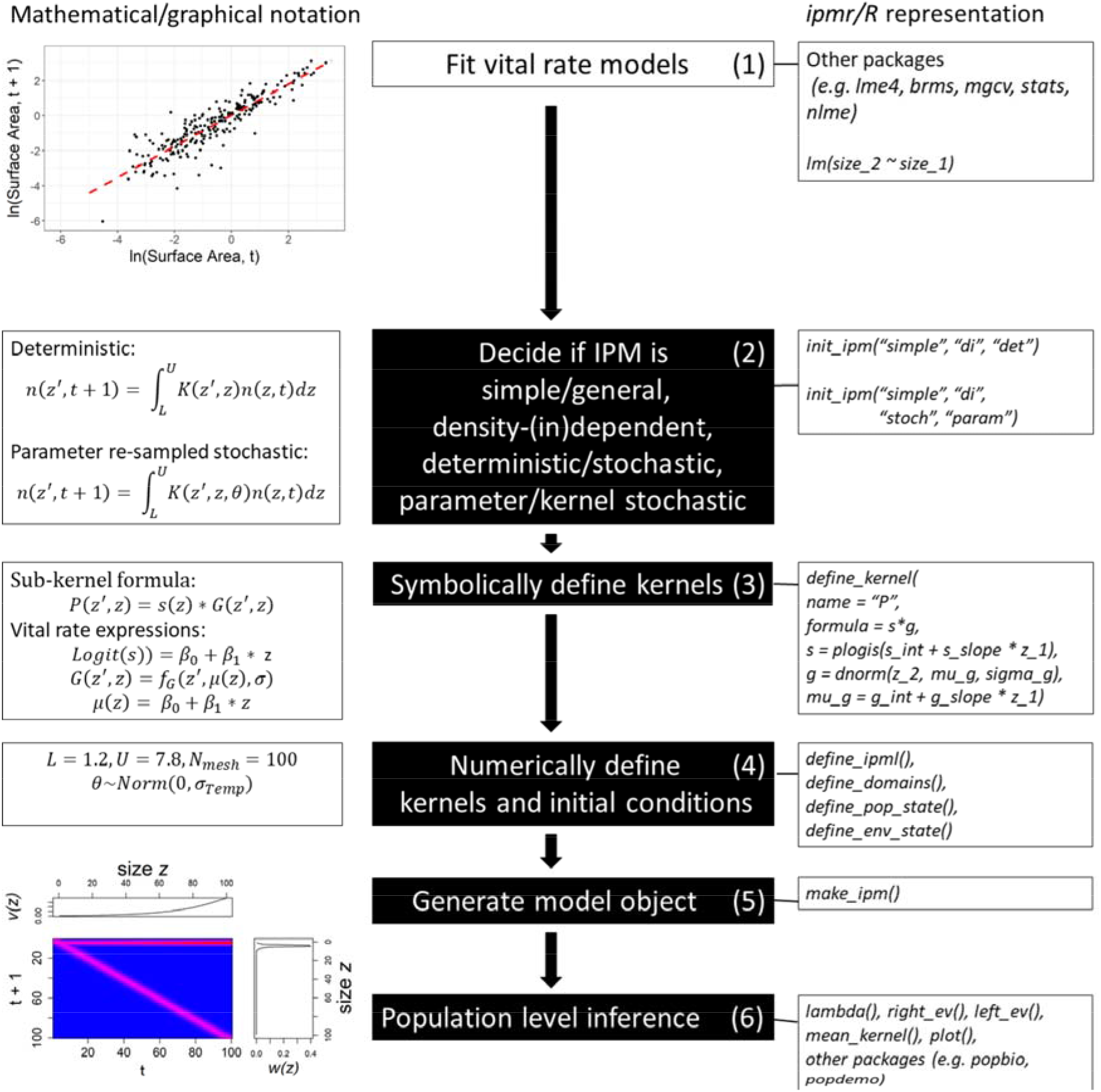
There are generally 6 steps in defining an IPM with `ipmr`. (1) Vital rate models are fit to demographic data collected from field sites. This step requires the use of other packages, as `ipmr` does not contain facilities for regression modeling. The figure on the left shows the fitted relationship between size at *t* and *t* + 1 for Carpobrotus spp. in Case Study 1. (2) The next step is deciding what type of IPM is needed. This is determined by both the research question and the data used to parameterize the regression models. This process is initiated with `init_ipm()`. In step (3), kernels are defined using `ipmr`’s syntax to represent kernels and vital rate functions. (4) Having defined symbolic representations of the model, the numerical definition is given. Here, the integration rule, domain bounds, and initial population conditions are defined. For some models, initial environmental conditions can also be defined. (5) `make_ipm()` numerically implements the proto_ipm object, (6) which can then be analyzed further. The figure at the bottom left shows a *K*(*z′, z*) kernel created by `make_ipm()` and `make_iter_kernel()`. The line plots above and to the right display the left and right eigenvectors, extracted with `left_ev()` and `right_ev()`, respectively.

**Table 1:** Translations between mathematical notation, R’s formula notation, and `ipmr`’s notation for the simplified version of Bogdan et al.’s Carpobrotus IPM. The `ipmr` column contains the expressions used in each kernel’s definition. *R* expressions are not provided for sub-kernels and model iteration procedures because they typically require defining functions separately, and there are many ways to do this step (examples are in the *R* code for each case study in the appendix). The plogis() function may not be familiar to some users. It is from the ‘stats’ *R* package and computes the inverse logit transformation of an expression.

| Math Formula | R Formula | ipmr |
| --- | --- | --- |
| $\mu^g = \alpha^g + \beta^g * z$ | size_2 ~ size_1, family = gaussian() | mu_g = g_int + g_slope * z |
| $g(z', z) = f^g(\mu^g, \sigma^g)$ | g = dnorm(z_2, mu_g, sd_g) | g = dnorm(z_2, mu_g, sd_g) |
| $logit(s(z)) = \alpha^s + \beta^s * z$ | surv ~ size_1, family = binomial() | s = plogis(s_int + s_slope * z) |
| $log(r^n(z)) = \alpha^{r^n} + \beta^{r^n} * z$ | fec ~ size_1, family = poisson() | r_n = exp(r_n_int + r_n_slope * z) |
| $logit(r^p(z)) = \alpha^{r^p} + \beta^{r^p} * z$ | repr ~ size_1, family = binomial() | r_p = plogis(r_p_int + r_p_slope * z) |
| $r^d(z') = f^{r^d}(\mu^{r^d}, \sigma^{r^d})$ | dnorm(z_2, mu_f_d, sigma_f_d) | r_d = dnorm(z_2, f_d_mu, f_d_sigma) |
| $p^r = \frac{\#Recruits(t+1)}{\#flowers(t)}$ | p_r = n_new_recruits / n_flowers | p_r = n_new / n_flowers |
| $P = s(z) * g(z', z)$ | | P = s * g |
| $F(z', z) = r^p(z) * r^n(z) * r^d(z') * p^r$ | | F = r_p * r_n * r_d * p_r |
| $n(z', t+1)$<br>$= \int_L^U [P(z', z) + F(z', z)]n(z, t)dz$ | | |

**Table 2:**
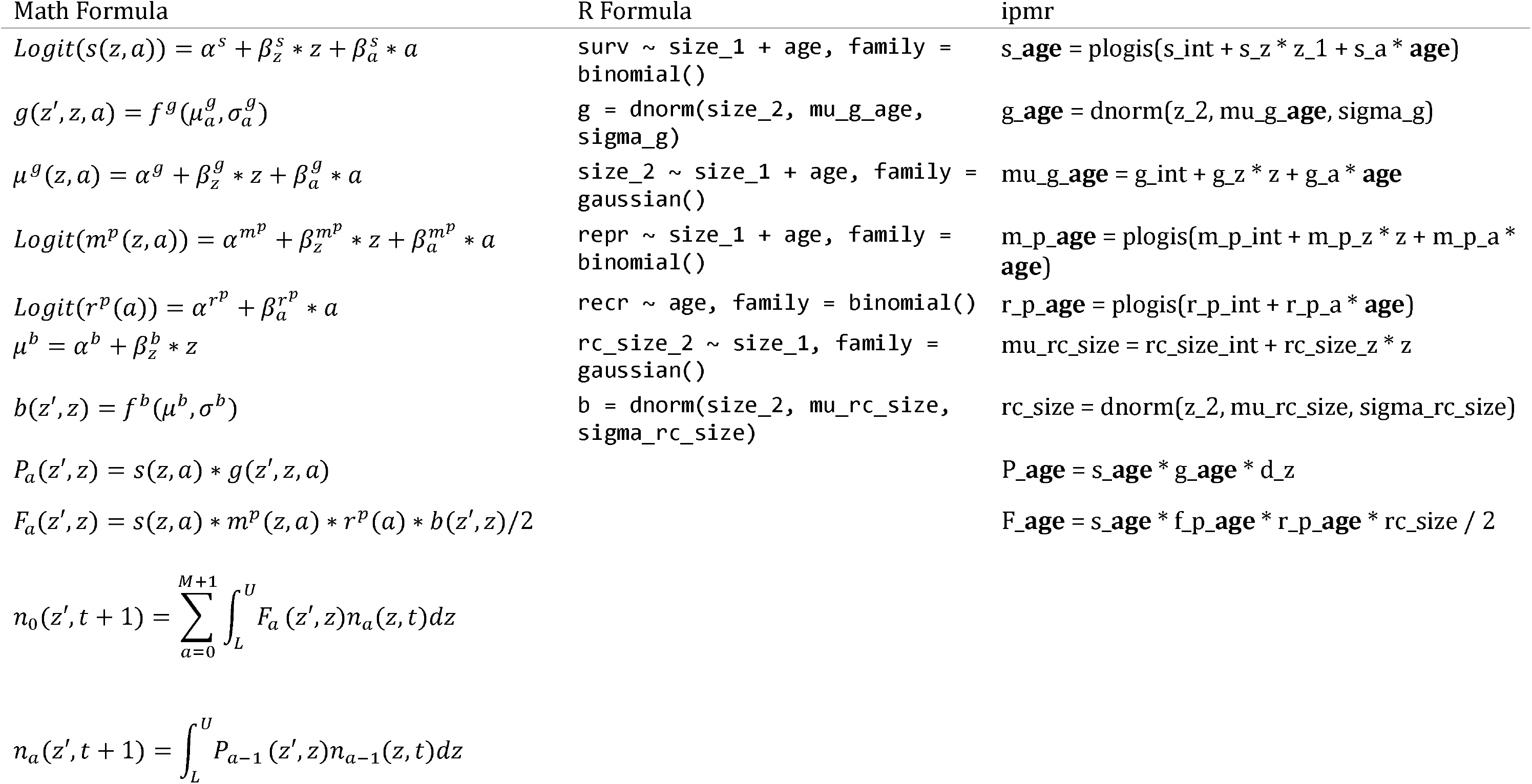
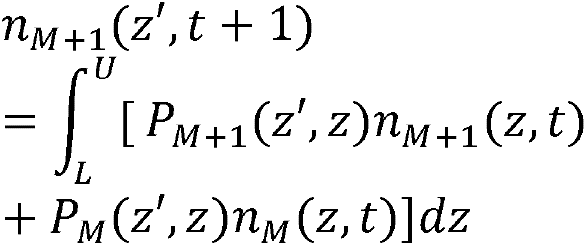
Translations between mathematical notation, R’s formula notation, and `ipmr`’s notation for Ellner et al. (2016) Ovis aries IPM. The `ipmr` column contains the expressions used in each kernel’s definition. *R* expressions are not provided for sub-kernels and model iteration procedures because they typically require defining functions separately, and there are many ways to do this step (examples are in the *R* code for each case study in the appendix). `ipmr` supports a suffix based syntax to avoid repetitively typing out the levels of discrete grouping variables. These are represented as ‘a’ in the Math column, ‘age’ in the *R* formula column, and are highlighted in bold in the `ipmr` column.

## Terminology and IPM construction

An IPM describes how the abundance and distribution of trait values (also called *state variables/states*, denoted *z* and *z*′) for a population changes in discrete time. The distribution of trait values in a population at time *t* is given by the function *n*(*z,t*). A simple IPM for the trait distribution *z*′ at time *t* + 1 is then

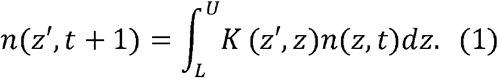

*K*(*z*′,*z*), known as the *projection kernel*, describes all possible transitions of existing individuals and recruitment of new individuals from *t* to *t* + 1, generating a new trait distribution *n*(*z′, t* + 1). *L, U* are the lower and upper bounds for values that the trait *z* can have, which defines the domain over which the integration is performed. The integral 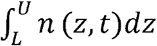 gives the total population size at time *t*.

To make the model more biologically interpretable, the projection kernel *K*(*z′, z*) is usually decomposed into sub-kernels (Eq 2). For example, a *projection kernel* to describe a lifecycle where individuals can survive, transition to different state values, and reproduce via sexual and asexual pathways, can be decomposed as follows

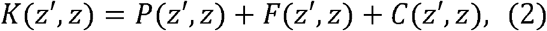

where *P*(*z′, z*) is a sub-kernel describing transitions due to survival and trait changes of existing individuals, *F*(*z′, z*) is a sub-kernel describing per-capita sexual contributions of existing individuals to recruitment, and *C*(*z′, z*) is a sub-kernel describing per-capita asexual contributions of existing individuals to recruitment. The sub-kernels are typically comprised of functions derived from regression models that relate an individual’s trait value *z* at time *t* to a new trait value *z*′ at *t* + 1. For example, the P kernel for Soay sheep (*Ovis aries*) on St. Kilda (Eq 3) may contain two regression models: (i) a logistic regression of survival on log body mass (Eq 4), and (ii) a linear regression of log body mass at *t* + 1 on log body mass at *t* (Eq 5–6). In this example, *f_g_* is a normal probability density function with *μ_g_* given by the linear predictor of the mean, and with *σ_g_* computed from the standard deviation of the residuals from the linear regression model.

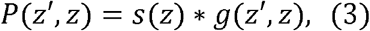

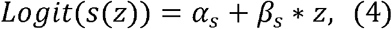

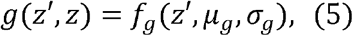

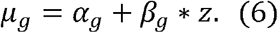

Analytical solutions to the integral in Eq 1 are usually not possible (Ellner & Rees, 2006). However, numerical approximations of these integrals can be constructed using a numerical integration rule. A commonly used rule is the midpoint rule (more complicated and precise methods are possible and will be implemented, though are not yet, see Ellner et al., 2016, Chapter 6). The midpoint rule divides the domain [*L, U*] into m artifical size bins centered at *z_i_* with width *h* = *(U - L)/m*. The midpoints *z_i_* = *L* + (*i* - 0.5) * *h* for *i* = 1, 2,…, *m*. The midpoint rule approximation for Eq 1 then becomes:

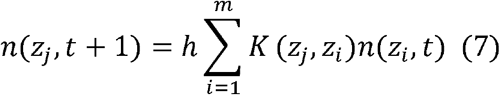

In practice, the numerical approximation of the integral converts the continuous projection kernel into a (large) discretized matrix. A matrix multiplication of the discretized projection kernel and the discretized trait distribution then generates a new trait distribution, a process referred to as model iteration (*sensu* Easterling et al., 2000).

Equations 1 and 2 are an example of a *simple IPM*. A critical aspect of `ipmr`‘s functionality is the distinction between *simple IPMs* and *general IPMs*. A simple IPM incorporates a single continuous state variable. Equations 1 and 2 represent a simple IPM because there is only one continuous state, *z*, and no additional discrete states. A general IPM models one or more continuous state variables, and/or discrete states. General IPMs are useful for modelling species with more complex life cycles. Many species’ life cycles contain multiple life stages that are not readily described by a single state variable. Similarly, individuals with similar trait values may behave differently depending on environmental context. For example, Bruno et al. (2011) modeled aspergillosis impacts on sea fan coral (*Gorgonia ventalina*) population dynamics by creating a model where colonies were cross classified by tissue area (continuously distributed) and infection status (a discrete state with two levels - infected and uninfected). Coulson, Tuljapurkar & Childs (2010) constructed a model for Soay sheep where the population was structured by body weight (continuously distributed) and age (discrete state). Mixtures of multiple continuous and discrete states are also possible. Indeed, the vital rates of many species with complex life cycles are often best described with multivariate state distributions (Caswell & Salguero-Gómez, 2013). A complete definition of the simple/general distinction is given in Ellner et al. (2016, Chapter 6).

### Case study 1 - A simple IPM

One use for IPMs is to evaluate potential performance and management of invasive species in their non-native range (*e.g.* Erickson et al., 2017). Calculating sensitivities and elasticities of *λ* to kernel perturbations can help identify conservation management strategies (de Kroon et al., 1986, Caswell, 2001, Baxter et al., 2006, Ellner et al., 2016). Bogdan et al. (2020) constructed a simple IPM for a *Carpobrotus* species growing north of Tel Aviv, Israel. The model includes four regressions, and an estimated recruit size distribution. Table 1 provides the mathematical formulae, the corresponding *R* model formula, and the `ipmr` notation for each one. The case study materials also offer an alternative implementation that uses the generic `predict()` function to generate the same output. The final part of the case study provides examples of functions that compute kernel sensitivity and elasticity, the per-generation growth rate, and generation time for the model, as well as how to visualize these results.

### Case study 2 - A general age X size IPM

We use an age- and size-structured IPM from Ellner et al. (2016) to illustrate how to create general IPMs with `ipmr`. This case study demonstrates is the suffix syntax for vital rate and kernel expressions, which is a key feature of `ipmr` (highlighted in bold in the ‘`ipmr`’ column in Table 2). The suffixes appended to each variable name in the `ipmr` formulation correspond to the sub- and/or super-scripts used in the mathematical formulation. `ipmr` internally expands the model expressions and substitutes the range of ages and/or grouping variables in for the suffixes. This allows users to specify their model in a way that closely mirrors its mathematical notation, and saves users from the potentially error-prone process of re-typing model definitions many times or using for loops over the range of discrete states. The case study then demonstrates how to couple age-specific survival and fertility with the model outputs.

## Discussion of additional applications

We have shown above how `ipmr` handles a variety of model implementations that go beyond the capabilities of existing scripts and packages. The underlying implementation based on metaprogramming should be able to readily incorporate future developments in parameterization methods. Regression modeling is a field that is constantly introducing new methods. As long as these new methods have functional forms for their expected value (or a method for `predict()`), `ipmr` should be able to implement IPMs using them. Finally, one particularly useful aspect of the package is the `proto_ipm` data structure. The `proto_ipm` is the common data structure used to represent every model class in `ipmr` and provides a concise, standardized format for representing IPMs. Furthermore, the `proto_ipm` object is created without any raw data, only functional forms and parameters. We are in the process of creating the PADRINO IPM database using `ipmr` and proto_ipms as an “engine” to re-build published IPMs using only functional forms and parameter estimates. This database could act as an IPM equivalent of the popular COMPADRE and COMADRE matrix population model databases (Salguero-Gómez et al., 2015, Salguero-Gómez et al., 2016). Recent work has highlighted the power of syntheses that harness many structured population models (Adler et al., 2013, Salguero-Gómez et al., 2016, Compagnoni et al., 2020). Despite the wide variety of models that are currently published in the IPM literature, `ipmr`’s functional approach is able to reproduce nearly all of them without requiring any raw data at all.

## Supporting information

ipmr Case Studies

## Author Contributions

All authors contributed to package design. SCL implemented the package. All authors wrote the first draft of the manuscript.

