## Supplementary material for "ipmr: Flexibly implement Integral Projection Models in R": ipmr Case Studies: 00_case_study_1.pdf

### Case Study 1: Bogdan et al. 2020

#### Two versions of a simple model

The first case study in this manuscript creates a model for *Carpobrotus spp.* The dataset used in this case study was collected in Havatselet Ha'Sharon, a suburb of Tel Aviv, Israel. The data were collected by drones taking aerial imagery of the population in successive years. Images were combined into a single high-resolution orthomosaic and georeferenced so the map from year 2 laid on top of the map from year 1. Flowers on each plant were counted using a point layer, and polygons were drawn around each ramet to estimate sizes and survival from year to year. Plants that had 0 flowers were classified as non-reproductive, and any plant with 1 or more flowers was classified as reproductive. This led to four regression models - survival, growth conditional on survival, probability of flowering, and number of flowers produced conditional on flowering. Finally, plants present in year 2 that were not present in year 1 were considered new recruits. The mean and variance of their sizes were computed, and this was used to model the recruit size distribution.

The resulting IPM is a simple IPM (i.e. no discrete states, one continuous state variable). The data that the regressions are fit to are included in the `ipmr` package, and can be accessed with `data(iceplant_ex)`.

The IPM can be written on paper as follows:

1. $n(z', t+1) = \int_L^U K(z', z) n(z, t) dz$
2. $K(z', z) = P(z', z) + F(z', z)$
3. $P(z', z) = s(z) * G(z', z)$
4. $F(z', z) = p_f(z) * r_s(z) * p_r * r_d(z')$

The components of each sub-kernel are either regression models or constants. Their functional forms are given below:

5. $Logit(s(z)) = \alpha_s + \beta_s * z$
6. $g(z', z) = f_g(z', \mu_g(z), \sigma_g)$
7. $\mu_g(z) = \alpha_g + \beta_g * z$
8. $Logit(p_f(z)) = \alpha_{p_f} + \beta_{p_f} * z$
9. $Log(r_s(z)) = \alpha_{r_s} + \beta_{r_s} * z$
10. $r_d(z') = f_{r_d}(z', \mu_{r_d}, \sigma_{r_d})$

$\alpha s$ and $\beta s$ correspond to intercepts and slopes from regression models, respectively. Here, $f_g$ and $f_{r_d}$ are used to denote normal probability density functions. The other parameters are constants derived directly from the data itself.

```
library(ipmr)

data(iceplant_ex)

# growth model.

grow_mod <- lm(log_size_next ~ log_size, data = iceplant_ex)
grow_sd  <- sd(resid(grow_mod))
```

```r
# survival model

surv_mod <- glm(survival ~ log_size, data = iceplant_ex, family = binomial())

# Pr(flowering) model

repr_mod <- glm(repro ~ log_size, data = iceplant_ex, family = binomial())

# Number of flowers per plant model

flow_mod <- glm(flower_n ~ log_size, data = iceplant_ex, family = poisson())

# New recruits have no size(t), but do have size(t + 1)

recr_data <- subset(iceplant_ex, is.na(log_size))

recr_mu  <- mean(recr_data$log_size_next)
recr_sd  <- sd(recr_data$log_size_next)

# This data set doesn't include information on germination and establishment.
# Thus, we'll compute the realized recruitment parameter as the number
# of observed recruits divided by the number of flowers produced in the prior
# year.

recr_n   <- length(recr_data$log_size_next)

flow_n   <- sum(iceplant_ex$flower_n, na.rm = TRUE)

recr_pr  <- recr_n / flow_n


# Now, we put all parameters into a list. This case study shows how to use
# the mathematical notation, as well as how to use predict() methods

all_params <- list(
  surv_int = coef(surv_mod)[1],
  surv_slo = coef(surv_mod)[2],
  repr_int = coef(repr_mod)[1],
  grow_int = coef(grow_mod)[1],
  grow_slo = coef(grow_mod)[2],
  grow_sdv = grow_sd,
  repr_slo = coef(repr_mod)[2],
  flow_int = coef(flow_mod)[1],
  flow_slo = coef(flow_mod)[2],
  recr_n   = recr_n,
  flow_n   = flow_n,
  recr_mu  = recr_mu,
  recr_sd  = recr_sd,
  recr_pr  = recr_pr
)
```

The next chunk generates a couple constants used to implement the model. We add 20% to the smallest and largest observed sizes to minimize eviction, and will implement the model with 100 meshpoints.

NB: `L` is multiplied by 1.2 because the log of the minimum observed size is negative, and we want to extend the size range to make it more negative. If `L` were positive, we'd multiply by 0.8.

```
L <- min(c(iceplant_ex$log_size,
           iceplant_ex$log_size_next),
         na.rm = TRUE) * 1.2

U <- max(c(iceplant_ex$log_size,
           iceplant_ex$log_size_next),
         na.rm = TRUE) * 1.2

n_mesh_p <- 100
```

We now have the parameter set prepared, and have the boundaries for our domains set up. We are ready to implement the model.

We start with the function `init_ipm()`. This function has five arguments: `sim_gen`, `di_dd`, `det_stoch`, `kern_param`, and `has_age`. For now, we will ignore the last argument, as it is covered in case study 2. The first 4 arguments specify the type of IPM we are building:

1. `sim_gen` - `simple/general`: This determines whether the model is a simple IPM or a general IPM.

2. `di_dd` - `di/dd`: This determines whether the model is density independent (`di`) or density dependent (`dd`).

3. `det_stoch` - `det/stoch`: This determines whether the model is deterministic (`det`) or stochastic (`stoch`). This particular model is deterministic, as there are no data on temporal or spatial changes in vital rates. An introduction to stochastic models is available here. This example does not make use of the final argument, `kern_param`, because it is not a stochastic model, so we'll ignore it for now.

Once we've decided on the type of model we want, we create the model class using one of the two options for each argument. Since there is no stochasticity, we can leave the fourth argument empty (its default is `NULL`). This case study is a simple, density independent, deterministic IPM, so we use the following:

```
carpobrotus_ipm <- init_ipm(sim_gen = "simple", di_dd = "di", det_stoch = "det")
```

After we have initialized our IPM, we need to start adding sub-kernels using the `define_kernel()` function. These correspond to equations 3 and 4 above. We'll start with the `P` kernel. It contains functions that describe survival of individual ramets, and, if they survive, their new sizes. Note that in `ipmr`, the order in which we define kernels for an IPM makes no difference, so we could also start with the `F` if we wanted to.

1. Survival is modeled with a logistic regression to predict the probability of survival to $t + 1$ based on the size of the ramet at $t$ (`surv_mod`). In order to use the coefficients from that model to generate a survival probability, we need to know the inverse logit transformation, or, a function that performs it for us based on the linear predictor.

2. Size at $t + 1$ is modeled with a Gaussian distribution with two parameters: the mean and standard deviation from the mean. The mean value of size at $t + 1$ (`mu_g`) is itself a linear function of size at $t$ and is parameterized with coefficients from the linear model (`grow_mod`). The standard deviation is a constant derived from the residual variance from the linear model we fit.

We start providing information on the `P` kernel by giving it a `name`. The name is important because we can use it to reference this kernel in higher level expressions later on. It can have any name we want, but `P` is consistent with the literature in this field (e.g. Easterling, Ellner & Dixon 2000, Ellner & Rees 2006). Next, we write the `formula`. The `formula` is the form of the kernel, and should look like Equation 3, without the $z$ and $z'$ arguments.

```
carpobrotus_ipm <-  define_kernel(
  proto_ipm = carpobrotus_ipm,
  name      = "P",
```

```
  formula    = s * G,
  ...
)
```

The `family` comes after `formula`. It describes the type of transition the kernel is implementing. `family` can be one of 4 options:

1. `"CC"`: Continuous state -> continuous state.
2. `"DC"`: discrete state -> continuous state.
3. `"CD"`: continuous state -> discrete state.
4. `"DD"`: discrete state -> discrete state.

Since this is a simple IPM with only 1 continuous state variable and 0 discrete state variables, the `family` will always be `"CC"`. In general IPMs, this will not always be true.

```
carpobrotus_ipm <-  define_kernel(
  proto_ipm = carpobrotus_ipm,
  name      = "P",
  formula   = s * G,
  family    = "CC",
  ...
)
```

We've now reached the `...` section of `define_kernel()`. The `...` part takes a set of named expressions that represent the vital rate functions we described in equations 5-7 above. The names on the left hand side of the `=` should appear either in the `formula` argument, or in other parts of the `...`. The expressions on the right hand side should generate the values that we want to plug in. For example, Equation 5 ($Logit(s(z)) = \alpha_s + \beta_s * z$) makes use of the `plogis` function in the `stats` package to compute the survival probabilities from our linear model. The names of the coefficients match the names in the `all_params` object we generated above. Another thing to note is the use of `z_1` and `z_2`. These are place-holders for $z, z'$ in the equations above. `ipmr` will generate values for these internally using information that we provide in some of the next steps.

```
carpobrotus_ipm <-  define_kernel(
  proto_ipm = carpobrotus_ipm,
  name      = "P",
  formula   = s * G,
  family    = "CC",
  G         = dnorm(z_2, mu_g, grow_sdv),
  mu_g      = grow_int + grow_slo * z_1,
  s         = plogis(surv_int + surv_slo * z_1),
  ...
)
```

After setting up our vital rate functions, the next step is to provide a couple more kernel-specific details:

1. `data_list`: this is the `all_params` object we created above. It contains the names and values of all the constants in our model.
2. `states`: A list that contains the names of the state variables in the kernel. In our case, we've just called them `"z"`. The `states` argument controls the names of the variables `z_1` and `z_2` that are generated internally. We could just as easily call them something else - we would just have to change the vital rate expressions to use those names instead. For example, in this model, $z, z'$ is the log-transformed surface area of ramets. We could abbreviate that with `"log_sa"`. In that case, `z_1,z_2` would become `log_sa_1, log_sa_2` in the vital rate expressions.
3. `evict_cor`: Whether or not to correct for eviction (Williams et al. 2012).

4. `evict_fun`: If we decide to correct for eviction, then a function that will correct it. In this example, we use `ipmr`'s `truncated_distributions` function. It takes two arguments: `fun`, which is the abbreviated form of the probability function family (e.g. "norm" for Gaussian, "lnorm" for log-normal, etc.), and `target`, which is the name in `...` that it modifies.

```
carpobrotus_ipm <-  define_kernel(
  proto_ipm = carpobrotus_ipm,
  name      = "P",
  formula   = s * G,
  family    = "CC",
  G         = dnorm(z_2, mu_g, grow_sdv),
  mu_g      = grow_int + grow_slo * z_1,
  s         = plogis(surv_int + surv_slo * z_1),
  data_list = all_params,
  states    = list(c("z")),
  evict_cor = TRUE,
  evict_fun = truncated_distributions(fun    = "norm",
                                      target = "G")
)
```

We've now defined our first sub-kernel. The next step is to repeat this process for the `F` kernel, which is Equations 4 and 8-10.

```
carpobrotus_ipm <-  define_kernel(
  proto_ipm = carpobrotus_ipm,
  name      = "F",
  formula   = recr_pr * r_s * r_d * p_f,
  family    = "CC",
  r_s       = exp(flow_int + flow_slo * z_1),
  r_d       = dnorm(z_2, recr_mu, recr_sd),
  p_f       = plogis(repr_int + repr_slo * z_1),
  data_list = all_params,
  states    = list(c("z")),
  evict_cor = TRUE,
  evict_fun = truncated_distributions(fun    = "norm",
                                      target =  "r_d")
)
```

We've defined our sub-kernels. The next step is tell `ipmr` how to implement it numerically, and provide the information needed to generate the correct iteration kernel. To do this, we use `define_impl()`, `define_domains()`, and `define_pop_state()`.

The first function tells `ipmr` which integration rule to use, which state variable each kernel acts on (`state_start`), and which state variable each kernel produces (`state_end`). The format of the list it takes in the `kernel_impl_list` argument can be tricky to implement right, so the helper function `make_impl_args_list()` makes sure everything is formatted properly. The `kernel_names` argument can be in any order. The `int_rule`, `state_start`, and `state_end` arguments are then matched to kernels in the `proto_ipm` based on the order in the `kernel_names`. Note that, at the moment, the only integration rule that's implemented is `"midpoint"`. `"b2b"` (bin to bin) and `"cdf"` (cumulative density functions) are in the works, and others can be implemented by popular demand.

```
carpobrotus_ipm <-  define_impl(
  proto_ipm = carpobrotus_ipm,
  make_impl_args_list(
    kernel_names = c("P", "F"),
    int_rule     = rep('midpoint', 2),
```

```
    state_start    = rep('z', 2),
    state_end      = rep('z', 2)
  )
)
```

Next, we define the range of values that our state variable, $z$/`z` can take on. This is done using `define_domains`. The `...` argument should have named vectors. The name should match the name of the `state`/`domain`. The first value in the vector is lower boundary, the second entry is the upper boundary, and the third entry is the number of bins to divide that range into.

```
carpobrotus_ipm <-  define_domains(
  proto_ipm = carpobrotus_ipm,
  z         = c(L, U, n_mesh_p)
)
```

Finally, we define the initial population state. In this case, we just use a uniform vector, but we could also use custom functions we defined on our own, or pre-specified vectors. The name of the population vector should be the name of the `state`/`domain`, with an `"n_"` attached to the front.

```
carpobrotus_ipm <-  define_pop_state(
  proto_ipm = carpobrotus_ipm,
    n_z     = rep(1/100, n_mesh_p)
)
```

Up until this point, all we've done is add components to the `proto_ipm`. We now have enough information in `proto_ipm` object to build a model, iterate it, and compute some basic quantities. `make_ipm()` is the next function we need. It generates the vital rate functions from the parameters and integration details we provided, and then builds the sub-kernels. At this point, it checks to make sure that everything makes numerical sense (e.g. there are no negative values or `NA`s generated). If we set `iterate = TRUE`, `make_ipm()` also generates expressions for iterating the model internally, and then evaluates those for the number of iterations supplied by `iterations`. There are a number of other arguments to `make_ipm()` that can prove helpful for subsequent analyses. `return_main_env` is one of these. The `main_env` object contains, among other things, the integration mesh and bin width information specified in `define_domains()`. We'll need the meshpoints and bin width for the analyses we'll do in the Further Analyses section, so we'll set `return_main_env = TRUE`.

```
carpobrotus_ipm <-  make_ipm(
  proto_ipm       = carpobrotus_ipm,
  iterate         = TRUE,
  iterations      = 100,
  return_main_env = TRUE
)


asymp_grow_rate <- lambda(carpobrotus_ipm)
asymp_grow_rate
```

```
## [1] 0.9759257
```

We see that the population is projected to shrink slightly. `ipmr` computes all values by iteration. Our measure of the asymptotic growth rate is the ratio $\frac{N_{t+1}}{N_t}$ for the final iteration of the model. If we are concerned about whether or not we've iterated our model enough to trust this value, we have two options: check for convergence using the helper `is_conv_to_asymptotic()`, or create the full iteration kernel, compute the dominant eigenvalue of that, and compare our estimate with the value obtained by iteration.

```
# Option 1: is_conv_to_asymptotic
```

```r
is_conv_to_asymptotic(carpobrotus_ipm)
```

```
## lambda
##   TRUE
```

```r
# Option 2: generate iteration kernel and compute eigenvalues

K <- make_iter_kernel(carpobrotus_ipm)

lam_eigen <- Re(eigen(K$mega_matrix)$values[1])

# If we've iterated our model enough, this should be approximately 0 (though
# maybe a little off due to floating point errors).

asymp_grow_rate - lam_eigen
```

```
## [1] 3.352874e-14
```

We can also inspect our sub-kernels, the time series of the population trait distribution, and make alterations to our model using some helpers from `ipmr`.

```r
# Sub-kernels have their own print method to display the range of values
# and some diagnotic information.

carpobrotus_ipm$sub_kernels
```

```
## $P
##
##  Minimum value: 0, maximum value: 0.08763
## All entries greater than or equal to 0: TRUE
##
## $F
##
##  Minimum value: 0, maximum value: 0.02512
## All entries greater than or equal to 0: TRUE
```

```r
# Extract the time series of the population state (n_z),
# and the n_t+1/n_t values (lambda)

pop_time_series    <- carpobrotus_ipm$pop_state$n_z
lambda_time_series <- carpobrotus_ipm$pop_state$lambda

# Next, we'll tweak the intercept of the p_f function and re-fit the model.

new_proto_ipm      <- carpobrotus_ipm$proto_ipm

# The parameters setter function takes a list. It can replace single values,
# create new values, or replace the entire parameter list, depending on how you
# set up the right hand side of the expression.

parameters(new_proto_ipm) <- list(repr_int = -0.3)

new_carp_ipm <- make_ipm(new_proto_ipm,
                         iterations = 100)

lambda(new_carp_ipm)
```

```
## [1] 0.9720439
```

Next, we'll go through an alternative implementation of the model using `predict(surv_mod)` instead of the mathematical form of the linear predictors. After that, we'll explore a couple additional analyses to see what is going on with this population of iceplants.

#### Using predict methods instead

We can simplify the code a bit more and get rid of the mathematical expressions for each regression model's link function by using `predict()` methods instead. The next chunk shows how to do this. Instead of extracting parameter values, we put the model objects themselves into the `data_list`. Next, we specify the `newdata` object where the name corresponds to the variable name(s) used in the model in question, and the values are the domain you want to evaluate the model on.

Above, we added parts to the `carpobrotus_ipm` object in a stepwise fashion. However, every `define_*` function in `ipmr` takes a `proto_ipm` as the first argument and returns a `proto_ipm` object. Thus, we can also use the `%>%` operator from the `magrittr` package to chain together the model creation pipeline. This example will demonstrate that process as well.

```
pred_par_list <- list(
  grow_mod = grow_mod,
  grow_sdv = grow_sd,
  surv_mod = surv_mod,
  repr_mod = repr_mod,
  flow_mod = flow_mod,
  recr_n   = recr_n,
  flow_n   = flow_n,
  recr_mu  = recr_mu,
  recr_sd  = recr_sd,
  recr_pr  = recr_pr
)

predict_method_carpobrotus <- init_ipm(sim_gen = "simple",
                                       di_dd = "di",
                                       det_stoch = "det") %>%
  define_kernel(
    name      = "P",
    formula   = s * G,
    family    = "CC",
    G         = dnorm(z_2, mu_g, grow_sdv),
    mu_g      = predict(grow_mod,
                        newdata = data.frame(log_size = z_1),
                        type = 'response'),
    s         = predict(surv_mod,
                        newdata = data.frame(log_size = z_1),
                        type = "response"),
    data_list = pred_par_list,
    states    = list(c('z')),
    evict_cor = TRUE,
    evict_fun = truncated_distributions("norm", "G")
  ) %>%
  define_kernel(
    name      = "F",
    formula   = recr_pr * r_s * r_d * p_f,
```

```
    family    = "CC",
    r_s       = predict(flow_mod,
                        newdata = data.frame(log_size = z_1),
                        type = "response"),
    r_d       = dnorm(z_2, recr_mu, recr_sd),
    p_f       = predict(repr_mod,
                        newdata = data.frame(log_size = z_1),
                        type = "response"),
    data_list = pred_par_list,
    states    = list(c("z")),
    evict_cor = TRUE,
    evict_fun = truncated_distributions("norm", "r_d")
  ) %>%
  define_impl(
    make_impl_args_list(
      kernel_names = c("P", "F"),
      int_rule     = rep('midpoint', 2),
      state_start    = rep('z', 2),
      state_end      = rep('z', 2)
    )
  ) %>%
  define_domains(
    z = c(L, U, n_mesh_p)
  )  %>%
  define_pop_state(
    n_z = rep(1/100, n_mesh_p)
  ) %>%
  make_ipm(iterate    = TRUE,
           iterations = 100)
```

#### Further analyses

Many research questions require a bit more than just computing asymptotic growth rate ($\lambda$). Below, we will compute the kernel sensitivity, elasticity, $R_0$, and generation time. First, we will define a couple helper functions. These are not included in `ipmr`, but will eventually be implemented in a separate package that can handle the various classes that `ipmr` works with.

The first is sensitivity of $\lambda$ to perturbations in the projection kernel. Here, we can use the `right_ev` and `left_ev` functions in `ipmr` to get the right and left eigenvectors, and then compute the sensitivity surface.

**Technical note:** `right_ev` and `left_ev` both compute eigenvectors via iteration. `left_ev` generates a transpose iteration using the `state_start` and `state_end` information contained in the `proto_ipm` object (defined in `define_impl`, for a full overview of transpose iteration, see Ellner & Rees, 2006, Appendix A). Because the form of for left iteration is different from the default of right iteration, `left_ev()` will always have to iterate a model. On the other hand, `right_ev` will always check to see if the model is already iterated. If so, and the population's trait distribution has converged to its asymptotic state, then it will just pull out the final distribution from the `ipm` object, scale it to sum to 1, and then return that without re-iterating anything. If not, it will use the final trait distribution from the `ipm` object as the starting point and iterate the model for 100 iterations (this can be adjusted as needed using the `iterations` argument to `right_ev`). If this fails to converge, it will return `NA` with a warning.

It is also important to note that we have a second argument here named `d_z`. This is the width of the integration bins. We'll see how to get that from our IPM below.

```
sens <- function(ipm_obj, d_z) {

  w <- right_ev(ipm_obj)[[1]]
  v <- left_ev(ipm_obj)[[1]]

  return(
    outer(v, w) / sum(v * w * d_z)
  )

}
```

Next, we can define a function to compute the elasticity of $\lambda$ to kernel perturbations. This uses the `sens` function from above, and the `lambda()` function from `ipmr`.

```
elas <- function(ipm_obj, d_z) {

  K           <- make_iter_kernel(ipm_obj)$mega_matrix

  sensitivity <- sens(ipm_obj, d_z)

  lamb        <- lambda(ipm_obj)

  out         <- sensitivity * (K / d_z) / lamb

  return(out)

}
```

We may also want to compute the per-generation population growth rate. The function below uses the sub-kernels contained in the `carpobrotus_ipm` object to do that.

```
R_nought <- function(ipm_obj) {

  Pm <- ipm_obj$sub_kernels$P
  Fm <- ipm_obj$sub_kernels$F

  I  <- diag(dim(Pm)[1])

  N  <- solve(I - Pm)

  R  <- Fm %*% N

  return(
    Re(eigen(R)$values)[1]
  )

}
```

Finally, generation time is a useful metric in many analyses. Below, we make use of our `R_nought` function to compute one version of this quantity (though other definitions exist. Covering those is beyond the scope of this case study).

```
gen_time <- function(ipm_obj) {

  lamb     <- unname(lambda(ipm_obj))
```

```
  r_nought <- R_nought(ipm_obj)

  return(log(r_nought) / log(lamb))
}
```

We need to extract the `d_z` value and meshpoints from the IPM we built. We can extract this information in a list form using the `int_mesh()` function from `ipmr` on our IPM object. The `d_z` in this case will be called `d_z` because we named our domain `"z"` when we implemented the model. However, it will have a different name if the `states` argument in `define_kernel` has different different values. Once we have that, we can begin computing all the values of interest.

```
mesh_info <- int_mesh(carpobrotus_ipm)

sens_mat <- sens(carpobrotus_ipm, mesh_info$d_z)
elas_mat <- elas(carpobrotus_ipm, mesh_info$d_z)

R0    <- R_nought(carpobrotus_ipm)
gen_T <- gen_time(carpobrotus_ipm)

R0
```

```
## [1] 0.5079748
```

```
gen_T
```

```
## [1] 27.79469
```

We may want to visualize the results of our sensitivity and elasticity analyses. We'll go through two options: one using the `graphics` package and one using the `ggplot2` package.

First, the `graphics` package.

```
par(mfrow = c(1, 2))

x <- y <- seq_len(ncol(sens_mat))

image(x = x,
      y = y,
      z = t(sens_mat),
      main = "Sensitivity", xlab = "t", ylab = "t + 1")

contour(x = x,
        y = y,
        t(sens_mat),
        nlevels = 5,
        labcex = 1.2,
        add = TRUE)

image(x = x,
      y = y,
      z = t(elas_mat),
      main = "Elasticity", xlab = "t", ylab = "t + 1",
      add = FALSE)
contour(x = x,
        y = y,
        t(elas_mat),
        nlevels = 5,
```

```
        labcex = 1.2,
        add = TRUE)
```

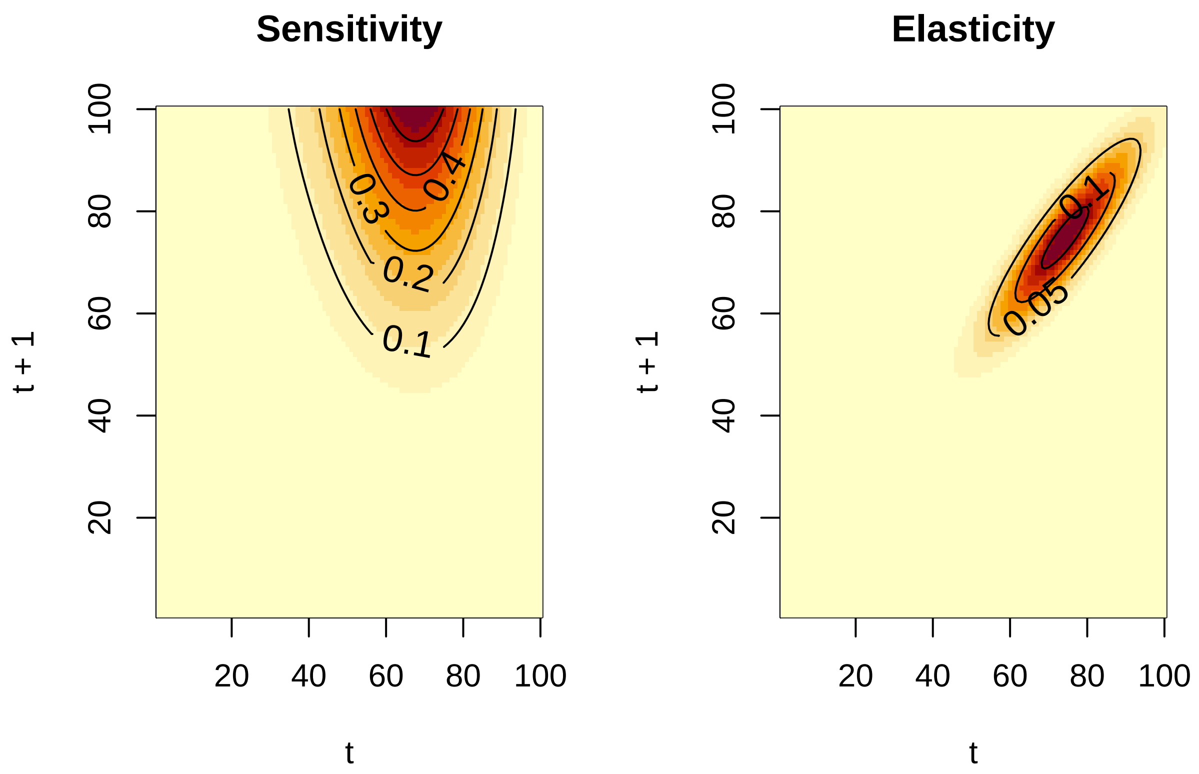


Now, for the `ggplot2` version. First, we create a long format of the matrix using `ipmr`'s `ipm_to_df` function. `ipm_to_df` can handle either bare matrices, or objects produced by `make_ipm`. The latter case is useful for plotting kernels directly using `ggplot2`. Once we've generated the long format sensitivity and elasticity matrices, we can use `geom_tile` and `geom_contour` to generate the `ggplots`, and `grid.arrange` from the `gridExtra` package to put them side by side.

```
library(ggplot2)
library(gridExtra)

sens_df <- ipm_to_df(sens_mat)
elas_df <- ipm_to_df(elas_mat)

# Create a default theme for our plots

def_theme <- theme(
  panel.background = element_blank(),
  axis.text        = element_blank(),
  axis.title.x     = element_text(
    size   = 16,
    margin = margin(
      t = 20,
      r = 0,
      l = 0,
      b = 2
```

```
    )
  ),
  axis.title.y = element_text(
    size   = 16,
    margin = margin(
      t = 0,
      r = 20,
      l = 2,
      b = 0
    )
  ),
  axis.ticks   = element_blank(),
  legend.title = element_text(size = 16)
)

sens_plt <- ggplot(sens_df) +
  geom_tile(aes(x    = t,
              y    = t_1,
              fill = value)) +
  geom_contour(aes(x = t,
                   y = t_1,
                   z = value),
              color = "black",
              size = 0.7) +
  scale_fill_gradient("Value",
                      low = "red",
                      high = "yellow") +
  scale_x_continuous(name = "t") +
  scale_y_continuous(name = "t + 1") +
  def_theme +
  theme(legend.title = element_blank()) +
  ggtitle("Sensitivity")

elas_plt <- ggplot(elas_df) +
  geom_tile(aes(x    = t,
              y    = t_1,
              fill = value)) +
  geom_contour(aes(x = t,
                   y = t_1,
                   z = value),
              color = "black",
              size = 0.7) +
  scale_fill_gradient("Value",
                      low = "red",
                      high = "yellow") +
  scale_x_continuous(name = "t") +
  scale_y_continuous(name = "t + 1") +
  def_theme +
  theme(legend.title = element_blank()) +
  ggtitle("Elasticity")

grid.arrange(sens_plt, elas_plt,
             layout_matrix = matrix(c(1, 2,
```

```
                                1, 2),
                              nrow = 2,
                              byrow = TRUE))
```

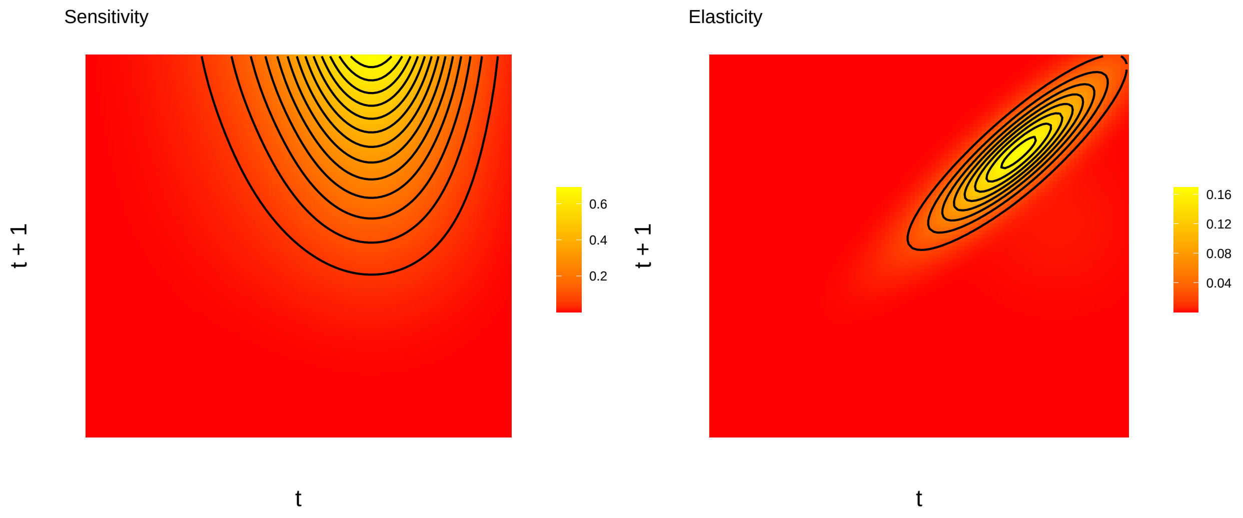
