## Supplementary material for "ipmr: Flexibly implement Integral Projection Models in R": ipmr Case Studies: 01_case_study_2.pdf

### Case Study 2: Ellner, Childs & Rees 2016

#### A more complicated, age and size structured model

Many life cycles cannot be described by a single, continuous state variable. For example, some plants are best modeled using height or diameter at breast height (DBH) as a state variable, and may also form a seed bank. Seeds in the seed bank can't have a value for height or DBH, but may lose their viability as they age. Thus, we require a discrete state to capture the dynamics of the seed bank. Models that include discrete states and/or multiple continuous state variables are *general IPMs*.

Many species exhibit age-dependent demography. Age may interact with a measure of size, for example body mass, resulting in neither single variable reliably predicting demography on its own. Data sets for which individuals are cross-classified by age and size represent an interesting opportunity for demographic research. This case study will use an age and size structured population to illustrate how to implement general IPMs in `ipmr`.

With size denoted $z, z'$, age denoted $a$, and the maximum age an individual can have denoted $M$, the model can be written as:

1. $n_0(z', t+1) = \sum\limits_{a=0}^{M} \int_L^U F_a(z', z) n_a(z, t) dz$
2. $n_a(z', t+1) = \int_L^U P_{a-1}(z', z) n_{a-1}(z, t) dz$ for $a = 1, 2, ..., M$

In this case, there is also an "greybeard" age class $M + 1$ defined as $a \geq M + 1$. We need to define one more equation to indicate the number of individuals in that age group.

3. $n_{M+1}(z', t+1) = \int_L^U [P_M(z', z) n_M(z, t) + P_{M+1}(z', z) n_{M+1}(z, t)] dz$

Below, $f_g$ and $f_{r_d}$ denote normal probability density functions. The sub-kernel $P_a(z', z)$ is comprised of the following functions:

4. $P_a(z', z) = s(z, a) * G(z', z, a)$
5. $Logit(s(z, a)) = \alpha_s + \beta_s^z * z + \beta_s^a * a$
6. $g(z', z, a) = f_g(z', \mu_g(z, a), \sigma_g)$
7. $\mu_g(z, a) = \alpha_g + \beta_g^z * z + \beta_g^a * a$

and the sub-kernel $F_a(z', z)$ is comprised of the following functions:

8. $F_0 = 0$
9. $F_a = s(z, a) * p_b(z, a) * p_r(a) * r_d(z', z) * 0.5$

The $F_a$ kernel is multiplied by 0.5 because this model only models the population dynamics of females.

10. $Logit(p_b(z, a)) = \alpha_{p_b} + \beta_{p_b}^z * z + \beta_{p_b}^a * a$
11. $Logit(p_r(a)) = \alpha_{p_r} + \beta_{p_r}^a * a$
12. $r_d(z', z) = f_{r_d}(z', \mu_{r_d}(z), \sigma_{r_d})$
13. $\mu_{r_d}(z) = \alpha_{r_d} + \beta_{r_d}^z * z$

Equations 5-7 and 10-13 are parameterized from regression models. The parameter values are taken from Ellner, Childs & Rees, Chapter 6 (2016). These can be found here. In the code from the book, these paramters were used to simulate an IBM to generate a data set. The data were then used to fit regression models and an age×size IPM. We are going to skip the simulation and regression model fitting steps steps and just use the "true" parameter estimates to generate the IPM.

First, we will define all the model parameters and a function for the $F_a$ kernels.

```
library(ipmr)

# Set parameter values and names

param_list <- list(
  ## Survival
  surv_int  = -1.70e+1,
  surv_z    =  6.68e+0,
  surv_a    = -3.34e-1,
  ## growth
  grow_int  =  1.27e+0,
  grow_z    =  6.12e-1,
  grow_a    = -7.24e-3,
  grow_sd   =  7.87e-2,
  ## reproduce or not
  repr_int  = -7.88e+0,
  repr_z    =  3.11e+0,
  repr_a    = -7.80e-2,
  ## recruit or not
  recr_int  =  1.11e+0,
  recr_a    =  1.84e-1,
  ## recruit size
  rcsz_int  =  3.62e-1,
  rcsz_z    =  7.09e-1,
  rcsz_sd   =  1.59e-1
)

# define a custom function to handle the F kernels. We could write a rather
# verbose if(age == 0) {0} else {other_math} in the define_kernel(), but that
# might look ugly. Note that we CANNOT use ifelse(), as its output is the same
# same length as its input (in this case, it would return 1 number, not 10000
# numbers).

r_fun <- function(age, s_age, pb_age, pr_age, recr) {

  if(age == 0) return(0)

  s_age * pb_age * pr_age * recr * 0.5

}
```

Next, we set up the $P_a$ kernels (Equations 4-7 above). Because this is a general, deterministic IPM, we use `init_ipm(sim_det = "general", di_dd = "di", det_stoch = "det", has_age = TRUE)`. We set `has_age = TRUE` to indicate that our model has age structure as well as size structure. There are 3 key things to note:

1. the use of the suffix `_age` appended to the names of the `"P_age"` kernel and the `mu_g_age` variable.

2. the value `age` used in the vital rate expressions.
3. the list in the `levels_ages` argument.

The values in the `levels_ages` list will automatically get substituted in for `"age"` each time it appears in the vital rate expressions and kernels. We add a second variable to this list, `"max_age"`, to indicate that we have a "greybeard" class. If we wanted our model to kill all individuals above age 20, we would simply omit the `"max_age"` slot in the `levels_ages` list.

This single call to `define_kernel()` will result in 22 actual kernels, one for each value of `age` from 0-21. For general IPMs that are not age-structured, we would use `has_hier_effs` and `levels_hier_effs` in the same way we're using `age` below.

The `plogis` function is part of the `stats` package in *R*, and performs the inverse logit transformation.

```
age_size_ipm <- init_ipm(sim_gen = "general",
                         di_dd = "di",
                         det_stoch = "det",
                         has_age = TRUE) %>%
  define_kernel(
    name           = "P_age",
    family         = "CC",
    formula        = s_age * g_age * d_z,
    s_age          = plogis(surv_int + surv_z * z_1 + surv_a * age),
    g_age          = dnorm(z_2, mu_g_age, grow_sd),
    mu_g_age       = grow_int + grow_z * z_1 + grow_a * age,
    data_list      = param_list,
    states         = list(c("z")),
    has_hier_effs = FALSE,
    levels_ages    = list(age = c(0:20), max_age = 21),
    evict_cor      = FALSE
  )
```

The $F_a$ kernel (equations 8-13) will follow a similar pattern - we append a suffix to the `name` paramter, and then make sure that our functions also include `_age` suffixes and `age` values where they need to appear.

```
age_size_ipm <-   define_kernel(
  proto_ipm     = age_size_ipm,
  name          = "F_age",
  family        = "CC",
  formula       = r_fun(age, s_age, pb_age, pr_age, recr) * d_z,
  s_age         = plogis(surv_int + surv_z * z_1 + surv_a * age),
  pb_age        = plogis(repr_int + repr_z * z_1 + repr_a * age),
  pr_age        = plogis(recr_int + recr_a * age),
  recr          = dnorm(z_2, rcsz_mu, rcsz_sd),
  rcsz_mu       = rcsz_int + rcsz_z * z_1,
  data_list     = param_list,
  states        = list(c("z")),
  has_hier_effs = FALSE,
  levels_ages   = list(age = c(0:20), max_age = 21),
  evict_cor     = FALSE
)
```

Once we've defined the $P_a$ and $F_a$ kernels, we need to define starting and ending states for each kernel. Age-size structured populations will look a little different from the previous ones, as we need to ensure that all fecundity kernels produce `age-0` individuals, and all survival-growth kernels produce `age` individuals. We define the implementation arguments using `define_impl()`, and set each kernel's `state_start` to `"z_age"`.

Because the fecundity kernel produces age-0 individuals, regardless of the starting age, its `state_end` is `"z_0"`.

```
age_size_ipm <-  define_impl(
  proto_ipm = age_size_ipm,
  make_impl_args_list(
    kernel_names = c("P_age", "F_age"),
    int_rule     = rep("midpoint", 2),
    state_start    = c("z_age", "z_age"),
    state_end      = c("z_age", "z_0")
  )
)
```

We define the domains using `define_domains()` in the same way we did for Case Study 1.

```
age_size_ipm <- age_size_ipm %>%
  define_domains(
    z = c(1.6, 3.7, 100)
  )
```

Our definition of the initial population state will look a little different though. We want to create 22 copies of the initial population state, one for each age group in the model. We do this by appending `_age` to the `n_z` in the `...` part of `define_pop_state`.

```
age_size_ipm <-  define_pop_state(
  proto_ipm = age_size_ipm,
  n_z_age = rep(1/100, 100)
  ) %>%
  make_ipm(
    usr_funs = list(r_fun = r_fun),
    iterate   = TRUE,
    iterations = 100
  )


lamb <- lambda(age_size_ipm)
lamb
```

```
## [1] 1.014833
```

#### Further analyses

The `right_ev` and `left_ev` functions also work for age × size models. We can use extract these, and plot them using a call to `lapply`.

NB: we assign the `lapply` call to a value here because `lines` returns `NULL` invisibly, and this clogs up the console. You probably don't need to do this for interactive use. We'll also divide the $dz$ value back into each eigenvector so that they are continuous distributions, rather than discretized vectors.

```
d_z <- int_mesh(age_size_ipm)$d_z

stable_dists <- right_ev(age_size_ipm)

w_plot <- lapply(stable_dists, function(x, d_z) x / d_z,
         d_z = d_z)
```

```r
repro_values <- left_ev(age_size_ipm)

v_plot <- lapply(repro_values, function(x, d_z) x / d_z,
                 d_z = d_z)


par(mfrow = c(1, 2))
plot(w_plot[[1]], type = 'l',
     ylab = expression(paste("w"[a],"(z)")),
     xlab = "Size bin")

x <- lapply(w_plot[2:22], function(x) lines(x))

plot(v_plot[[1]], type = 'l',
     ylab = expression(paste("v"[a],"(z)")),
     xlab = "Size bin",
     ylim = c(0, 0.2))

x <- lapply(v_plot[2:22], function(x) lines(x))
```

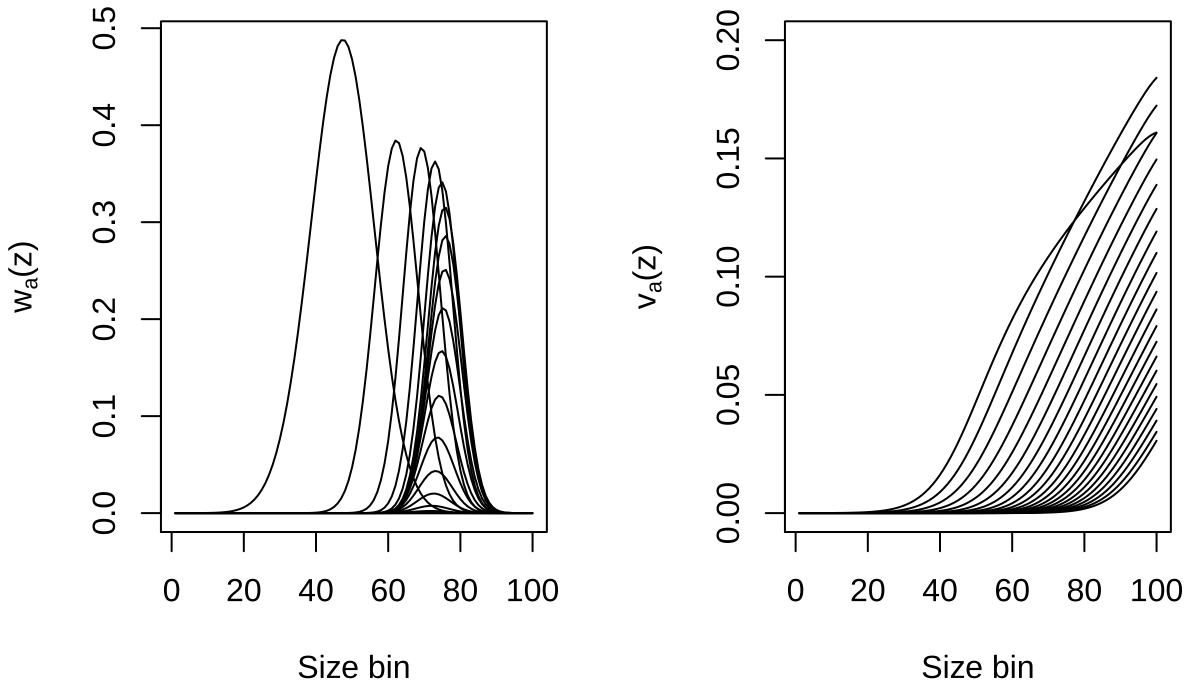


Next, we'll compute age-specific survival ($\tilde{l}_a$/`l_a`) and fecundity ($\tilde{f}_a$/`f_a`) values. These are defined as follows:

$$\tilde{l}_a = eP^a c$$

$$\tilde{f}_a = (eFP^a c)/l_a$$

where $c$ is some distribution of newborns.

We'll initialize a cohort using the stable size distribution for age-0 individuals that we obtained above. Next, we'll iterate them through our model for 100 years, and see who's left, and how much they reproduced.

NB: do not try to use this method for computing $R_0$ - it will lead to incorrect results because in this particular model, parental state affects initial offspring state. For more details, see Ellner, Childs & Rees (2016), Chapters 3 and 6.

A couple technical notes:

1. We are going to split out our P and F sub-kernels into separate lists so that indexing them is easier during the iteration process.
2. We need to set our `max_age` variable to 22 now, so that we don't accidentally introduce an "off-by-1" error when we index the sub-kernels in our IPM object.
3. We use an identity matrix to compute the initial value of `l_a`, because by definition all age-0 individuals must survive to age-0.

```
# Initialize a cohort and some vectors to hold our quantities of interest.

init_pop <- stable_dists[[1]] / sum(stable_dists[[1]])
n_yrs    <- 100L

l_a <- f_a <- numeric(n_yrs)

P_kerns <- age_size_ipm$sub_kernels[grepl("P", names(age_size_ipm$sub_kernels))]
F_kerns <- age_size_ipm$sub_kernels[grepl("F", names(age_size_ipm$sub_kernels))]

# We have to bump max_age because R indexes from 1, and our minimum age is 0.

max_age <- 22

P_a <- diag(length(init_pop))

for(yr in seq_len(n_yrs)) {

  # When we start, we want to use age-specific kernels until we reach max_age.
  # after that, all survivors have entered the "greybeard" class.

  if(yr < max_age) {

    P_now <- P_kerns[[yr]]
    F_now <- F_kerns[[yr]]

  } else {

    P_now <- P_kerns[[max_age]]
    F_now <- F_kerns[[max_age]]

  }

  l_a[yr] <- sum(colSums(P_a) * init_pop)
  f_a[yr] <- sum(colSums(F_now %*% P_a) * init_pop)

  P_a <- P_now %*% P_a
```

```
}

f_a <- f_a / l_a

# Looks like most are dead at after 25 years, so we'll restrict our
# plot range to that time span

par(mfrow = c(1, 2))

plot(l_a[1:25], type = 'l',
     ylab = expression(paste("l"[a])),
     xlab = "Age")
plot(f_a[1:25], type = 'l',
     ylab = expression(paste("f"[a])),
     xlab = "Age")
```

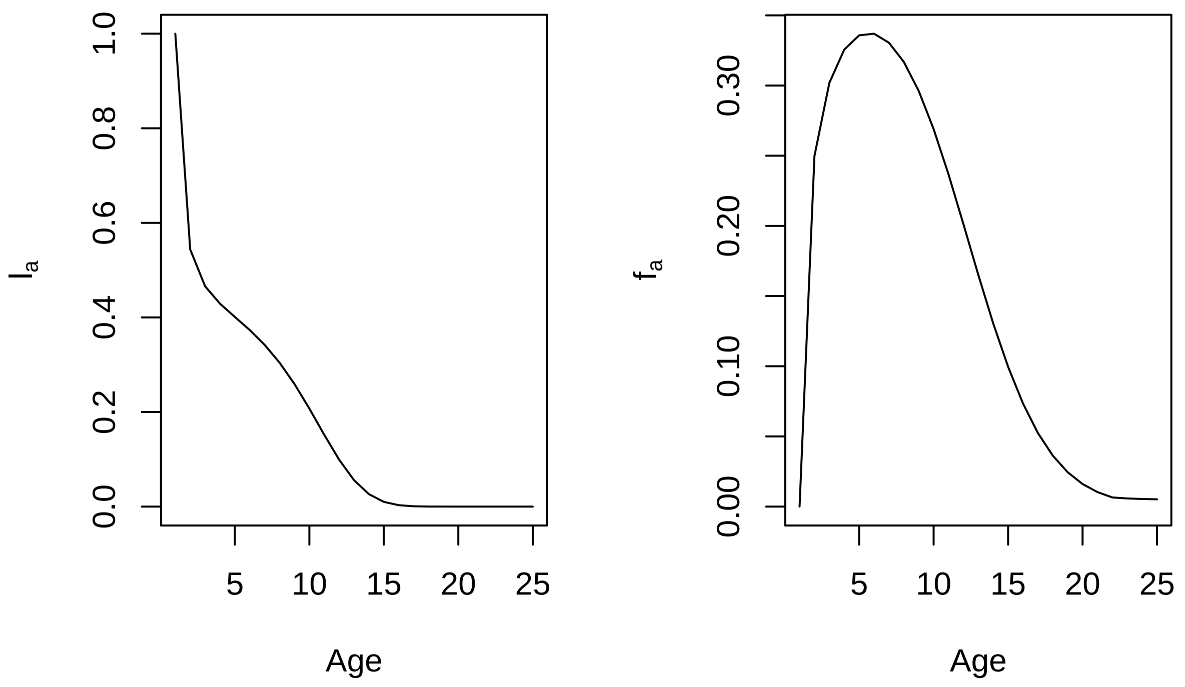


We can also calculate the age-specific survival probability $p_a$ as $\frac{l_{a+1}}{l_a}$. We'll restrict our calculations to the to 25 years.

```
p_a <- l_a[2:26] / l_a[1:25]

plot(p_a, type = 'l',
     ylab = expression(paste("p"[a])),
     xlab = "Age")
```

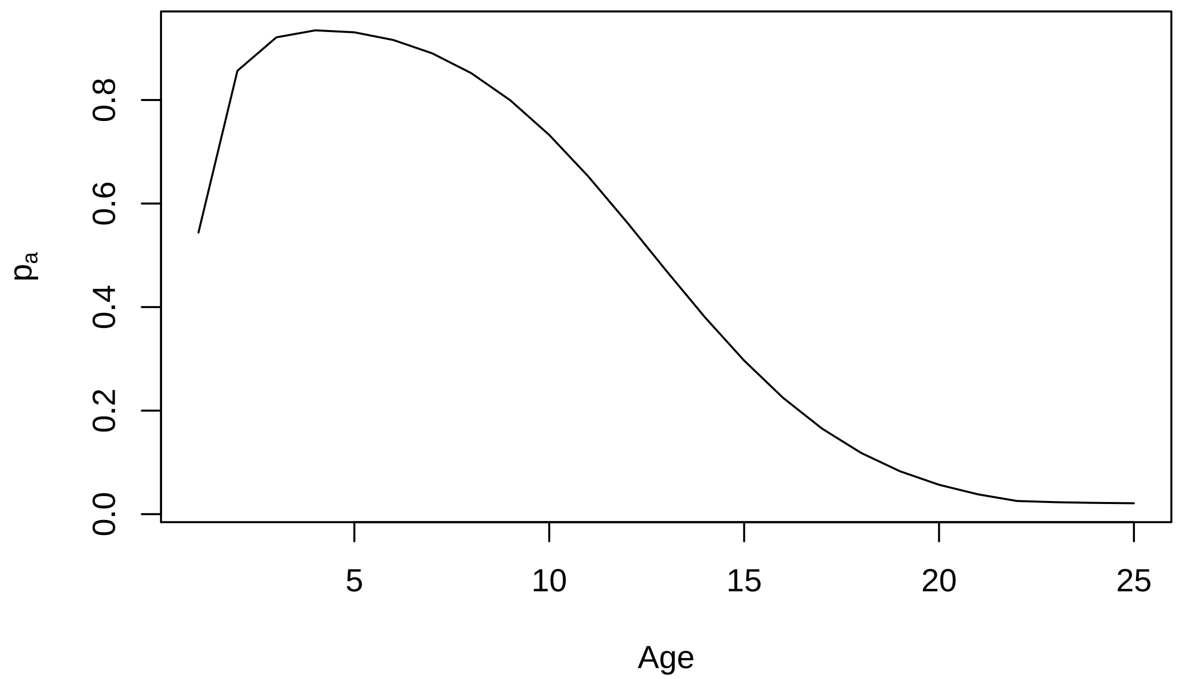

0.0
0.2
0.4
0.6
0.8
$p_a$
5
10
15
20
25
Age
