## Supplementary material for "ipmr: Flexibly implement Integral Projection Models in R": ipmr Case Studies: 02_SOM.pdf

### Supplementary Online Materials for ipmr: Flexibly implement Integral Projection Models in R

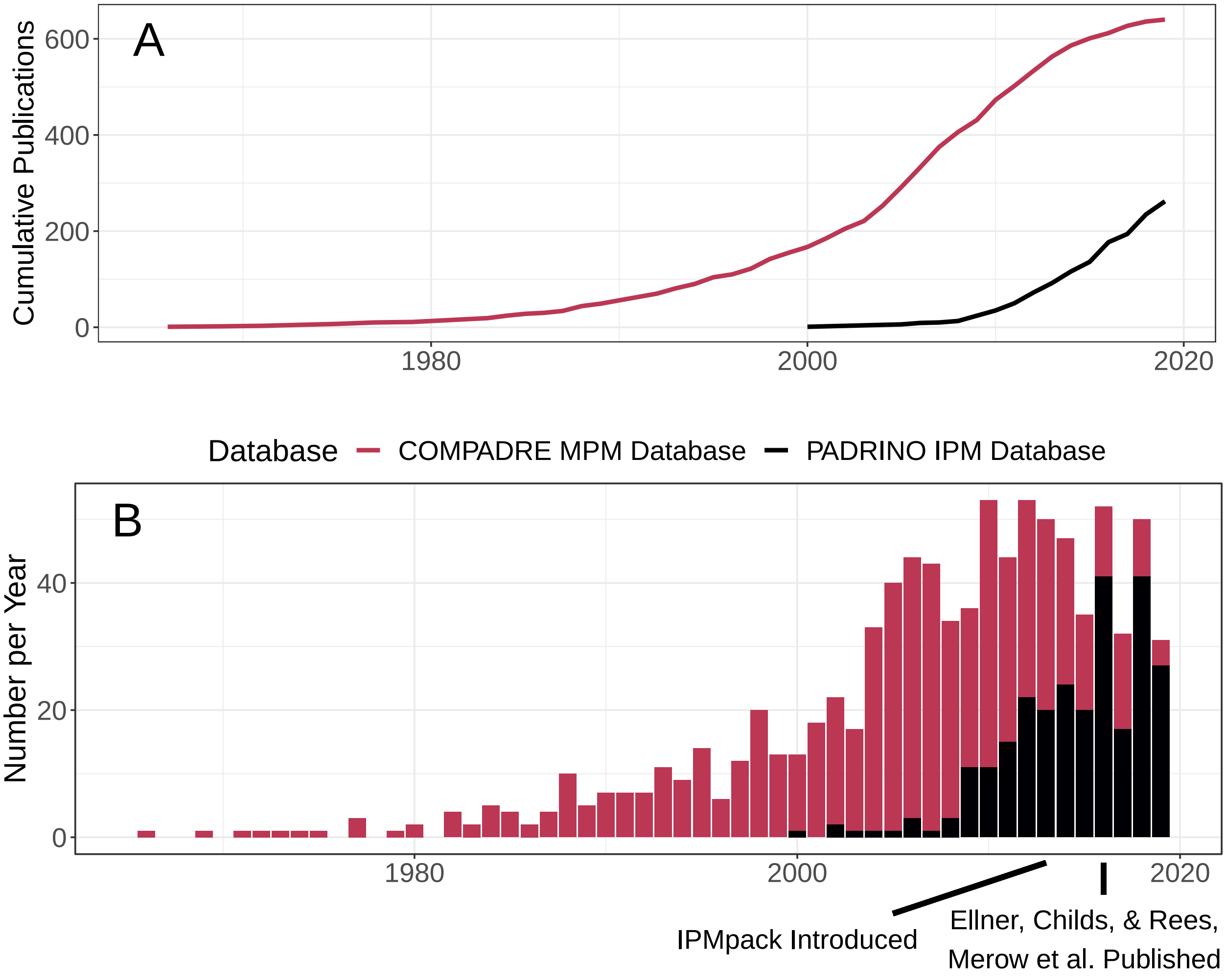


Figure 1: Figure S1: The usage of integral projection models (IPMs) has increased rapidly since their introduction. Cumulative number of publications using matrix projection models (MPMs, red) and IPMs (black) (A) and number of publications per year for each type of model (B). IPMs have been adopted rapidly since their introduction in 2000. Unfortunately, software packages to assist with their implementation have not kept pace with their theoretical advancements and applications to ever more complex demographic data.

Table 1: Table S1: Table of species, kingdom, and publications for papers that contain Integral Projection Models. This table is not exhaustive and does not include some papers for which species and kingdom information is not relevant (e.g. papers developing new theory relying on simulated data).

| Species | Kingdom | Full Citation |
|---|---|---|
| Celtis zenkeri | Plantae | Picard, N., & Liang, J. (2014). Matrix models for size-structured populations: unrealistic fast growth or simply diffusion?. PloS one, 9(6), e98254. |
| Staudtia kamerunensis | Plantae | Picard, N., & Liang, J. (2014). Matrix models for size-structured populations: unrealistic fast growth or simply diffusion?. PloS one, 9(6), e98254. |
| Coelocaryon preussii | Plantae | Picard, N., & Liang, J. (2014). Matrix models for size-structured populations: unrealistic fast growth or simply diffusion?. PloS one, 9(6), e98254. |
| Musanga cecropioides | Plantae | Picard, N., & Liang, J. (2014). Matrix models for size-structured populations: unrealistic fast growth or simply diffusion?. PloS one, 9(6), e98254. |
| Carapa procera | Plantae | Picard, N., & Liang, J. (2014). Matrix models for size-structured populations: unrealistic fast growth or simply diffusion?. PloS one, 9(6), e98254. |
| Garcinia punctata | Plantae | Picard, N., & Liang, J. (2014). Matrix models for size-structured populations: unrealistic fast growth or simply diffusion?. PloS one, 9(6), e98254. |
| Dasylepis seretii | Plantae | Picard, N., & Liang, J. (2014). Matrix models for size-structured populations: unrealistic fast growth or simply diffusion?. PloS one, 9(6), e98254. |
| Trichilia rubescens | Plantae | Picard, N., & Liang, J. (2014). Matrix models for size-structured populations: unrealistic fast growth or simply diffusion?. PloS one, 9(6), e98254. |
| Rinorea oblongifolia | Plantae | Picard, N., & Liang, J. (2014). Matrix models for size-structured populations: unrealistic fast growth or simply diffusion?. PloS one, 9(6), e98254. |
| Pycnanthus angolensis | Plantae | Picard, N., & Liang, J. (2014). Matrix models for size-structured populations: unrealistic fast growth or simply diffusion?. PloS one, 9(6), e98254. |
| Pancovia laurentii | Plantae | Picard, N., & Liang, J. (2014). Matrix models for size-structured populations: unrealistic fast growth or simply diffusion?. PloS one, 9(6), e98254. |
| Trilepisium madagas-cariense | Plantae | Picard, N., & Liang, J. (2014). Matrix models for size-structured populations: unrealistic fast growth or simply diffusion?. PloS one, 9(6), e98254. |
| Diospyros iturensis | Plantae | Picard, N., & Liang, J. (2014). Matrix models for size-structured populations: unrealistic fast growth or simply diffusion?. PloS one, 9(6), e98254. |
| Petersianthus macrocarpus | Plantae | Picard, N., & Liang, J. (2014). Matrix models for size-structured populations: unrealistic fast growth or simply diffusion?. PloS one, 9(6), e98254. |
| Eribroma oblongum | Plantae | Picard, N., & Liang, J. (2014). Matrix models for size-structured populations: unrealistic fast growth or simply diffusion?. PloS one, 9(6), e98254. |
| Synsepalum stipulatum | Plantae | Picard, N., & Liang, J. (2014). Matrix models for size-structured populations: unrealistic fast growth or simply diffusion?. PloS one, 9(6), e98254. |
| Trichilia prieurieana | Plantae | Picard, N., & Liang, J. (2014). Matrix models for size-structured populations: unrealistic fast growth or simply diffusion?. PloS one, 9(6), e98254. |
| Scottellia coriacea | Plantae | Picard, N., & Liang, J. (2014). Matrix models for size-structured populations: unrealistic fast growth or simply diffusion?. PloS one, 9(6), e98254. |
| Drypetes chevalieri | Plantae | Picard, N., & Liang, J. (2014). Matrix models for size-structured populations: unrealistic fast growth or simply diffusion?. PloS one, 9(6), e98254. |
| Angylocalyx pynaertii | Plantae | Picard, N., & Liang, J. (2014). Matrix models for size-structured populations: unrealistic fast growth or simply diffusion?. PloS one, 9(6), e98254. |
| Manilkara mabokensis | Plantae | Picard, N., & Liang, J. (2014). Matrix models for size-structured populations: unrealistic fast growth or simply diffusion?. PloS one, 9(6), e98254. |
| Manilkara pellegriniana | Plantae | Picard, N., & Liang, J. (2014). Matrix models for size-structured populations: unrealistic fast growth or simply diffusion?. PloS one, 9(6), e98254. |

| | | | |
|---|---|---|---|
| Polyalthia suaveolens | | Plantae | Picard, N., & Liang, J. (2014). Matrix models for size-structured populations: unrealistic fast growth or simply diffusion?. PloS one, 9(6), e98254. |
| Pausinystalia macroceras | | Plantae | Picard, N., & Liang, J. (2014). Matrix models for size-structured populations: unrealistic fast growth or simply diffusion?. PloS one, 9(6), e98254. |
| Entandrophragma cylindricum | | Plantae | Picard, N., & Liang, J. (2014). Matrix models for size-structured populations: unrealistic fast growth or simply diffusion?. PloS one, 9(6), e98254. |
| Macaranga paxii | | Plantae | Picard, N., & Liang, J. (2014). Matrix models for size-structured populations: unrealistic fast growth or simply diffusion?. PloS one, 9(6), e98254. |
| Celtis mildbraedii | | Plantae | Picard, N., & Liang, J. (2014). Matrix models for size-structured populations: unrealistic fast growth or simply diffusion?. PloS one, 9(6), e98254. |
| Guarea laurentii | | Plantae | Picard, N., & Liang, J. (2014). Matrix models for size-structured populations: unrealistic fast growth or simply diffusion?. PloS one, 9(6), e98254. |
| Diospyros canaliculata | | Plantae | Picard, N., & Liang, J. (2014). Matrix models for size-structured populations: unrealistic fast growth or simply diffusion?. PloS one, 9(6), e98254. |
| Drypetes gilgiana | | Plantae | Picard, N., & Liang, J. (2014). Matrix models for size-structured populations: unrealistic fast growth or simply diffusion?. PloS one, 9(6), e98254. |
| Cola lateritia | | Plantae | Picard, N., & Liang, J. (2014). Matrix models for size-structured populations: unrealistic fast growth or simply diffusion?. PloS one, 9(6), e98254. |
| Strombosia grandifolia | | Plantae | Picard, N., & Liang, J. (2014). Matrix models for size-structured populations: unrealistic fast growth or simply diffusion?. PloS one, 9(6), e98254. |
| Dialium guineense | | Plantae | Picard, N., & Liang, J. (2014). Matrix models for size-structured populations: unrealistic fast growth or simply diffusion?. PloS one, 9(6), e98254. |
| Diospyros crassiflora | | Plantae | Picard, N., & Liang, J. (2014). Matrix models for size-structured populations: unrealistic fast growth or simply diffusion?. PloS one, 9(6), e98254. |
| Corynanthe pachyceras | | Plantae | Picard, N., & Liang, J. (2014). Matrix models for size-structured populations: unrealistic fast growth or simply diffusion?. PloS one, 9(6), e98254. |
| Triplochiton scleroxylon | | Plantae | Picard, N., & Liang, J. (2014). Matrix models for size-structured populations: unrealistic fast growth or simply diffusion?. PloS one, 9(6), e98254. |
| Lecaniodiscus cupanioides | | Plantae | Picard, N., & Liang, J. (2014). Matrix models for size-structured populations: unrealistic fast growth or simply diffusion?. PloS one, 9(6), e98254. |
| Anonidium mannii | | Plantae | Picard, N., & Liang, J. (2014). Matrix models for size-structured populations: unrealistic fast growth or simply diffusion?. PloS one, 9(6), e98254. |
| Cola nitida | | Plantae | Picard, N., & Liang, J. (2014). Matrix models for size-structured populations: unrealistic fast growth or simply diffusion?. PloS one, 9(6), e98254. |
| Funtumia elastica | | Plantae | Picard, N., & Liang, J. (2014). Matrix models for size-structured populations: unrealistic fast growth or simply diffusion?. PloS one, 9(6), e98254. |
| Santiria trimera | | Plantae | Picard, N., & Liang, J. (2014). Matrix models for size-structured populations: unrealistic fast growth or simply diffusion?. PloS one, 9(6), e98254. |
| Pouteria altissima | | Plantae | Picard, N., & Liang, J. (2014). Matrix models for size-structured populations: unrealistic fast growth or simply diffusion?. PloS one, 9(6), e98254. |
| Chrysophyllum africanum | | Plantae | Picard, N., & Liang, J. (2014). Matrix models for size-structured populations: unrealistic fast growth or simply diffusion?. PloS one, 9(6), e98254. |
| Celtis adolfi-friderici | | Plantae | Picard, N., & Liang, J. (2014). Matrix models for size-structured populations: unrealistic fast growth or simply diffusion?. PloS one, 9(6), e98254. |
| Strombosiopsis tetrandra | | Plantae | Picard, N., & Liang, J. (2014). Matrix models for size-structured populations: unrealistic fast growth or simply diffusion?. PloS one, 9(6), e98254. |
| Aubrevillea kerstingii | | Plantae | Picard, N., & Liang, J. (2014). Matrix models for size-structured populations: unrealistic fast growth or simply diffusion?. PloS one, 9(6), e98254. |
| Drypetes sp. | | Plantae | Picard, N., & Liang, J. (2014). Matrix models for size-structured populations: unrealistic fast growth or simply diffusion?. PloS one, 9(6), e98254. |
| Ricinodendron heudelotii | | Plantae | Picard, N., & Liang, J. (2014). Matrix models for size-structured populations: unrealistic fast growth or simply diffusion?. PloS one, 9(6), e98254. |

| | | |
|---|---|---|
| Entandrophragma angolense | Plantae | Picard, N., & Liang, J. (2014). Matrix models for size-structured populations: unrealistic fast growth or simply diffusion?. PloS one, 9(6), e98254. |
| Drypetes obanensis | Plantae | Picard, N., & Liang, J. (2014). Matrix models for size-structured populations: unrealistic fast growth or simply diffusion?. PloS one, 9(6), e98254. |
| Khaya anthotheca | Plantae | Picard, N., & Liang, J. (2014). Matrix models for size-structured populations: unrealistic fast growth or simply diffusion?. PloS one, 9(6), e98254. |
| Albizia glaberrima | Plantae | Picard, N., & Liang, J. (2014). Matrix models for size-structured populations: unrealistic fast growth or simply diffusion?. PloS one, 9(6), e98254. |
| Chrysophyllum lacourtianum | Plantae | Picard, N., & Liang, J. (2014). Matrix models for size-structured populations: unrealistic fast growth or simply diffusion?. PloS one, 9(6), e98254. |
| Sancassania berlesei | Animalia | Ozgul A, Coulson T, Reynolds A, Cameron TC & Benton TG (2012) Population responses to perturbations: the importance of trait-based analysis illustrated through a microcosm experiment. American Naturalist 179: 582-594 |
| Phyteuma spicatum | Plantae | Kolb A (2012) Differential effects of herbivory and pathogen infestation on plant population dynamics. Plant Ecology 213: 315-326 |
| Phyteuma spicatum | Plantae | Kolb A, Dahlgren JP & Ehrlén (2010) Population size affects vital rates but not population growth rate of a perennial plant. Ecology 91: 3210-3217 |
| Tridacna maxima | Animalia | Yau, A. J. Y. (2011). Size-based approaches to modeling and managing local populations: A case study using an artisanal fishery for giant clams, Tridacna maxima. University of California, Santa Barbara. |
| Tridacna maxima | Animalia | Yau, A. J., Lenihan, H. S., & Kendall, B. E. (2014). Fishery management priorities vary with self-recruitment in sedentary marine populations. Ecological Applications, 24(6), 1490-1504. |
| Cirsium vulgare | Plantae | Tenhumberg, B., Suwa, T., Tyre, A. J., Russell, F. L., & Louda, S. M. (2015). Integral projection models show exotic thistle is more limited than native thistle by ambient competition and herbivory. Ecosphere, 6(4), art69. |
| Aconitum noveborancense | Plantae | Easterling MR, Ellner SP & Dixon PM (2000) Size-specific sensitivity: applying a new structured population model. Ecology 81: 694-708 |
| Oenothera glazioviana | Plantae | Godfray, H. Charles J., and Mark Rees. "Population growth rates: issues and an application." Philosophical Transactions of the Royal Society B: Biological Sciences 357.1425 (2002): 1307-1319. |
| Aeonium arboreum | Plantae | Pannell, J. L. (2016). Climatic limitation of alien weeds in New Zealand: enhancing species distribution models with field data (Doctoral dissertation, Lincoln University). |
| Aeonium haworthii | Plantae | Pannell, J. L. (2016). Climatic limitation of alien weeds in New Zealand: enhancing species distribution models with field data (Doctoral dissertation, Lincoln University). |
| Cecropia obtusifolia | Plantae | Metcalf CJE, Horvitz CC, Tuljapurkar S & Clark DA (2009) A time to grow and a time to die: a new way to analyze the dynamics of size, light, age, and death of tropical trees. Ecology 90: 2766-2778 |
| Cecropia insignis | Plantae | Metcalf CJE, Horvitz CC, Tuljapurkar S & Clark DA (2009) A time to grow and a time to die: a new way to analyze the dynamics of size, light, age, and death of tropical trees. Ecology 90: 2766-2778 |
| Simarouba amara | Plantae | Metcalf CJE, Horvitz CC, Tuljapurkar S & Clark DA (2009) A time to grow and a time to die: a new way to analyze the dynamics of size, light, age, and death of tropical trees. Ecology 90: 2766-2778 |
| Minquartia guianensis | Plantae | Metcalf CJE, Horvitz CC, Tuljapurkar S & Clark DA (2009) A time to grow and a time to die: a new way to analyze the dynamics of size, light, age, and death of tropical trees. Ecology 90: 2766-2778 |
| Balizia elegans | Plantae | Metcalf CJE, Horvitz CC, Tuljapurkar S & Clark DA (2009) A time to grow and a time to die: a new way to analyze the dynamics of size, light, age, and death of tropical trees. Ecology 90: 2766-2778 |

| | | |
|---|---|---|
| Hymenolobium mesoameri-canum | Plantae | Metcalf CJE, Horvitz CC, Tuljapurkar S & Clark DA (2009) A time to grow and a time to die: a new way to analyze the dynamics of size, light, age, and death of tropical trees. Ecology 90: 2766-2778 |
| Lecythis ampla | Plantae | Metcalf CJE, Horvitz CC, Tuljapurkar S & Clark DA (2009) A time to grow and a time to die: a new way to analyze the dynamics of size, light, age, and death of tropical trees. Ecology 90: 2766-2778 |
| Dipteryx panamensis | Plantae | Metcalf CJE, Horvitz CC, Tuljapurkar S & Clark DA (2009) A time to grow and a time to die: a new way to analyze the dynamics of size, light, age, and death of tropical trees. Ecology 90: 2766-2778 |
| Hyeronima alchorneoides | Plantae | Metcalf CJE, Horvitz CC, Tuljapurkar S & Clark DA (2009) A time to grow and a time to die: a new way to analyze the dynamics of size, light, age, and death of tropical trees. Ecology 90: 2766-2778 |
| Hypericum cumulicola | Plantae | Metcalf CJE, McMahon SM, Salguero-Gómez R & Jongejans E (2013) IPMpack: an R package for Integral Projection Models. Methods in Ecology and Evolution 4: 195-200 |
| Arabidopsis thaliana | Plantae | Metcalf CJE & Mitchell-Olds T (2009) Life history in a model system: opening the black box with Arabidopsis thaliana. Ecology Letters 12: 593-600 |
| Carduus nutans | Plantae | Metcalf CJE, Rees M, Buckley YM & Sheppard AW (2009) Seed predators and the evolutionary stable flowering staregy in the invasive plant Carduus nutans. Evolutionary Ecology 23: 893-906 |
| Oenothera glazioviana | Plantae | Metcalf CJE, Rose KE & Rees M (2003) Evolutionary demography of monocarpic perennials. Trends in Ecology and Evolution 18: 471-180 |
| Carlina vulgaris | Plantae | Metcalf, C. J. E., Ellner, S., Childs, D. Z., Salguero?Gómez, R., Merow, C., McMahon, S. M., ... & Rees, M. (2015). Statistical modeling of annual variation for inference on stochastic population dynamics using Integral Projection Models. Methods in Ecology and Evolution. |
| Ovis aries | Animalia | Metcalf, C. J. E., Ellner, S., Childs, D. Z., Salguero?Gómez, R., Merow, C., McMahon, S. M., ... & Rees, M. (2015). Statistical modeling of annual variation for inference on stochastic population dynamics using Integral Projection Models. Methods in Ecology and Evolution. |
| Ovis aries | Animalia | Childs DZ, Coulson TN, Pemberton JM, Clutton-Brock TH & Rees M (2011) Predicting trait values and measuring selection in complex life histories: reproductive allcoation decisions in Soay sheep. Ecology Letters 14: 985-992 |
| Polygonum cuspidatum | Plantae | Dauer JT & Jongejans E (2013) Elucidating the population dynamics of Japanese knotweed using integral projection models. PLOS ONE in 8:e75181 |
| Mammillaria dixanthocentron | Plantae | González, E. J., Rees, M., & Martorell, C. (2013). Identifying the demographic processes relevant for species conservation in human-impacted areas: does the model matter?. Oecologia, 171(2), 347-356. |
| Mammillaria hernandezii | Plantae | González, E. J., Rees, M., & Martorell, C. (2013). Identifying the demographic processes relevant for species conservation in human-impacted areas: does the model matter?. Oecologia, 171(2), 347-356. |
| Actaea spicata | Plantae | Ehrlén, J., Raabova, J., & Dahlgren, J. P. (2015). Flowering schedule in a perennial plant-Life-history trade-offs, seed predation and total offspring fitness. Ecology. |
| Borderea pyrenaica | Plantae | García MB, Dahlgren JP & Ehrlén J (2011) No evidence of senescence in a 300-year-old mountain herb. Journal of Ecology 99: 1424-1430 |
| Alliaria petiolata | Plantae | Merow, C., Bois, S. T., Allen, J. M., Xie, Y., Silander Jr., J. A. (2017). Climate change both facilitates and inhibits invasive plant ranges in New England. PNAS Vol. 114, Issue 16. |
| Onopordum illyricum | Plantae | Ellner SP & Rees M (2006) Integral projection models for species with complex demography. American Naturalist 167: 410-428 |

| | | |
|---|---|---|
| Onopordum illyricum | Plantae | Ellner SP & Rees M (2007) Stochastic stable population growth in integral projection models: theory and application. Journal of Mathematical Biology 54: 227-256 |
| Lepidium latifolium | Plantae | Ellner SP & Schreiber SJ (2012) Temporarilly variable dispersal and demography can accelerate the spread of invading species. Theoretical Population Biology 82: 283-298 |
| Veratrum album | Plantae | Hesse E, Rees M & Müller-Schärer (2008) Life-history variation in contrasting habitats: flowering decision in a clonal perennial herb (Veratrum album). American Naturalist 172: 196-213 |
| Crematogaster laevis | Animalia | Bruna, E. M., Izzo, T. J., Inouye, B. D., & Vasconcelos, H. L. (2014). Effect of mutualist partner identity on plant demography. Ecology, 95(12), 3237-3243. |
| Pheidole minutula | Animalia | Bruna, E. M., Izzo, T. J., Inouye, B. D., & Vasconcelos, H. L. (2014). Effect of mutualist partner identity on plant demography. Ecology, 95(12), 3237-3243. |
| Ovis aries | Animalia | Simmonds, E. G., & Coulson, T. (2015). Analysis of phenotypic change in relation to climatic drivers in a population of Soay sheep Ovis aries. Oikos, 124(5), 543-552. |
| Carduus nutans | Plantae | Jongejans E, Shea K, Skarpaas O, Kelly D & Ellner SP (2011) Importance of individual and environmental variation for invasive species spread: a spatial integral projection model. Ecology 92: 86-97 |
| Caragana intermedia | Plantae | Li, S. L., Yu, F. H., Werger, M. J., Dong, M., Ramula, S., & Zuidema, P. A. (2013). Understanding the effects of a new grazing policy: the impact of seasonal grazing on shrub demography in the I nner M ongolian steppe. Journal of Applied Ecology, 50(6), 1377-1386. |
| Artemisia ordosica | Plantae | Li S-L, Yu F-H, Werger MJA, Dong M & Zuidema PA (2011) Habitat-specific demography across dune fixation stages in a semi-arid sandland: understanding the expansion, stabilization and decline of a dominant shrub. Journal of Ecology 99: 610-620 |
| Capreolus capreolus | Animalia | Plard, F., Gaillard, J.-M., Coulson, T., Delorme, D., Warnant, C., Michallet, J., Tuljapurkar, S., Krishnakumar, S., Bonenfant, C. (2015), Quantifying the influence of measured and unmeasured individual differences on demography. Journal of Animal Ecology. doi: 10.1111/1365-2656.12393 |
| Cervus elaphus | Animalia | Plard, F., Gaillard, J. M., Coulson, T., Hewison, A. M., Douhard, M., Klein, F., ... & Bonenfant, C. (2015). The influence of birth date via body mass on individual fitness in a long-lived mammal. Ecology. |
| Lepomis gibbosus | Animalia | van Kleef, H. H., & Jongejans, E. (2014, September). Identifying drivers of pumpkinseed invasiveness using population models. In Aquatic Invasions (Vol. 9, No. 3, pp. 315-326). Regional Euro-Asian Biological Invasions Centre (REABIC). |
| Orchis purpurea | Plantae | Jacquemyn H, Brys R & Jongejans E (2010) Size-dependent flowering and costs of reproduction affect population dynamics in a tuberous perennial woodland orchid. Journal of Ecology 98: 1204-1215 |
| Bouteloua eripoda | Plantae | Chu, C., & Adler, P. B. (2014). When should plant population models include age structure?. Journal of ecology, 102(2), 531-543. |
| Bouteloua rothrockii | Plantae | Chu, C., & Adler, P. B. (2014). When should plant population models include age structure?. Journal of ecology, 102(2), 531-543. |
| Heteropogon contortus | Plantae | Chu, C., & Adler, P. B. (2014). When should plant population models include age structure?. Journal of ecology, 102(2), 531-543. |
| Hilaria belangeri | Plantae | Chu, C., & Adler, P. B. (2014). When should plant population models include age structure?. Journal of ecology, 102(2), 531-543. |
| Hesperostipa comata | Plantae | Chu, C., & Adler, P. B. (2014). When should plant population models include age structure?. Journal of ecology, 102(2), 531-543. |
| Koeleria maculata | Plantae | Chu, C., & Adler, P. B. (2014). When should plant population models include age structure?. Journal of ecology, 102(2), 531-543. |

| | | |
|---|---|---|
| Poa secunda | Plantae | Chu, C., & Adler, P. B. (2014). When should plant population models include age structure?. Journal of ecology, 102(2), 531-543. |
| Pseudoroegneria spicata | Plantae | Chu, C., & Adler, P. B. (2014). When should plant population models include age structure?. Journal of ecology, 102(2), 531-543. |
| Astridia longiseta | Plantae | Chu, C., & Adler, P. B. (2014). When should plant population models include age structure?. Journal of ecology, 102(2), 531-543. |
| Boutelona curtipendula | Plantae | Chu, C., & Adler, P. B. (2014). When should plant population models include age structure?. Journal of ecology, 102(2), 531-543. |
| Boutelona gracilis | Plantae | Chu, C., & Adler, P. B. (2014). When should plant population models include age structure?. Journal of ecology, 102(2), 531-543. |
| Boutelona hirsuta | Plantae | Chu, C., & Adler, P. B. (2014). When should plant population models include age structure?. Journal of ecology, 102(2), 531-543. |
| Muhlenbergia arenacea | Plantae | Chu, C., & Adler, P. B. (2014). When should plant population models include age structure?. Journal of ecology, 102(2), 531-543. |
| Scleropogon brevifoloius | Plantae | Chu, C., & Adler, P. B. (2014). When should plant population models include age structure?. Journal of ecology, 102(2), 531-543. |
| Sporobplus flexuosus | Plantae | Chu, C., & Adler, P. B. (2014). When should plant population models include age structure?. Journal of ecology, 102(2), 531-543. |
| Rhizoglyphus robini | Animalia | Smallegange, I. M., Deere, J. A., & Coulson, T. (2014). Correlative changes in life-history variables in response to environmental change in a model organism. The American Naturalist, 183(6), 784-797. |
| Rhizoglyphus robini | Animalia | Smallegange, I. M., Deere, J. A., & Coulson, T. (2014). Correlative changes in life-history variables in response to environmental change in a model organism.The American Naturalist, 183(6), 784-797. |
| Capreolus capreolus | Animalia | Plard, F., Gaillard, J. M., Coulson, T., Hewison, A. M., Delorme, D., Warnant, C., & Bonenfant, C. (2014). Mismatch between birth date and vegetation phenology slows the demography of roe deer. PLoS Biol, 12(4), e1001828. |
| Cervus elaphus | Animalia | Plard, F., Gaillard, J. M., Coulson, T., Hewison, A. M., Delorme, D., Warnant, C., & Bonenfant, C. (2014). Mismatch between birth date and vegetation phenology slows the demography of roe deer. PLoS Biol, 12(4), e1001828. |
| Goodyera pubescens | Plantae | Diez, J. M., Giladi, I., Warren, R., & Pulliam, H. R. (2014). Probabilistic and spatially variable niches inferred from demography. Journal of ecology, 102(2), 544-554. |
| Agrostics hyemalis | Plantae | Williams JL, Miller TX & Ellner SP (2012) Avoiding unintentional eviction from integral projection models. Ecology 93: 2008-2014 |
| Anemone patens | Plantae | Williams JL, Miller TX & Ellner SP (2012) Avoiding unintentional eviction from integral projection models. Ecology 93: 2008-2014 |
| Opuntia imbricata | Plantae | Williams JL, Miller TX & Ellner SP (2012) Avoiding unintentional eviction from integral projection models. Ecology 93: 2008-2014 |
| Anemone patens | Plantae | Williams JL & Crone EE (2006) The impact of invasive grasses on the population growth of Anemone patens, a long-lived native forb. Ecology 87: 3200-3208 |
| Ranunculus weyleri | Plantae | Cursach, J., Besnard, A., Rita, J., & Fréville, H. (2013). Demographic variation and conservation of the narrow endemic plant Ranunculus weyleri. Acta Oecologica, 53, 102-109. |
| Dracocephalum austriacum | Plantae | Nicolè F, Dahlgren JP, Vivat A, Till-Bottraud & Ehrlén J (2011) Interdependent effects of habitat quality and climate on population growth of an endangered plant. Journal of Ecology 99: 1211-1218 |
| Actaea spicata | Plantae | Dahlgren JP & Ehrlén J (2009) Linking environmental variation to population dynamics of a forest herb. Journal of Ecology 97: 666-674 |
| Acatea spicata | Plantae | Dahlgren, J. P., & Ehrlén, J. (2011). Incorporating environmental change over succession in an integral projection model of population dynamics of a forest herb. Oikos, 120(8), 1183-1190. |

| | | | |
|---|---|---|---|
| Borderea pyrenaica | | Plantae | Dahlgren JP, García MB & Ehrlén J (2011) Nonlinear relationships between vital rates and state variables in demographic models. Ecology 92: 1181-1187 |
| Lathyrus vernus | | Plantae | Dahlgren, J. P., Östergård, H., & Ehrlén, J. (2014). Local environment and density-dependent feedbacks determine population growth in a forest herb. Oecologia, 176(4), 1023-1032. |
| Pseudoroegneria spicata | | Plantae | Dalgleish HJ, Koons DN, Hooten MB, Moffet CA & Adler PB (2011) Climate influences the demography of three dominant sagebrush steppe plants. Ecology 92: 75-85 |
| Hesperostipa comata | | Plantae | Dalgleish HJ, Koons DN, Hooten MB, Moffet CA & Adler PB (2011) Climate influences the demography of three dominant sagebrush steppe plants. Ecology 92: 75-85 |
| Artemisia tripartita | | Plantae | Dalgleish HJ, Koons DN, Hooten MB, Moffet CA & Adler PB (2011) Climate influences the demography of three dominant sagebrush steppe plants. Ecology 92: 75-85 |
| Gorgonia ventalina | | Animalia | Bruno JF, Ellner SP, Vu I, Kim K & Harvell CD (2011) Impacts of aspergillosis on sea fan coral demography: modeling a moving target. Ecological Monographs 81: 123-139 |
| Acropora hyacinthus | | Animalia | Madin JS, Hughes TP & Connolly SR (2012) Calcification, storm damage and population resilience of tabular corals under climate change. PLoS One 7: 1-10 |
| Dendrophylax lindenii | | Plantae | Raventós, J., González, E., Mújica, E., & Doak, D. F. (2015). Population Viability Analysis of the Epiphytic Ghost Orchid (Dendrophylax lindenii) in Cuba. Biotropica, 47(2), 179-189. |
| Cynoglossum officinale | | Plantae | Williams JL (2009) Flowering life-history strategies differ between the native and introduced ranges of a monocarpic perennial. American Naturalist 174: 660-672 |
| Cynoglossum officinale | | Plantae | Williams JL, Auge H & Maron JL (2010) Testing hypotheses for exotic plant success: parallel experiments in the native and introduced ranges. Ecology 91: 1355-1366 |
| Poulsenia armata | | Plantae | Zambrano, J., & Salguero?Gómez, R. (2014). Forest Fragmentation Alters the Population Dynamics of a Late?successional Tropical Tree. Biotropica, 46(5), 556-564. |
| Cirsium canescens | | Plantae | Rose KE, Louda SM & Rees M (2005) Demographic and evolutionary impacts of native and invasive herbivores on Cirsium canescens. Ecology 86: 453-465 |
| Crocodylus niloticus | | Animalia | Wallace W, Leslie A & Coulson T. 2012. Re-evaluating the effect of harvesting regimes on Nile crocodiles using an integral projection models. In press |
| Cirsium altissimum | | Plantae | Rose, K. E., Russell, F. L., & Louda, S. M. (2011). Integral projection model of insect herbivore effects on Cirsium altissimum populations along productivity gradients. Ecosphere, 2(8), art97. |
| Agrostis hyemalis | | Plantae | Yule, K. M., Miller, T. E., & Rudgers, J. A. (2013). Costs, benefits, and loss of vertically transmitted symbionts affect host population dynamics. Oikos, 122(10), 1512-1520. |
| Phoenix loureiroi | | Plantae | Mandle, L., Ticktin, T., & Zuidema, P. A. (2015). Resilience of palm populations to disturbance is determined by interactive effects of fire, herbivory and harvest. Journal of Ecology. |
| Ovis canadensis | | Animalia | Traill, L. W., Schindler, S., & Coulson, T. (2014). Demography, not inheritance, drives phenotypic change in hunted bighorn sheep. Proceedings of the National Academy of Sciences, 111(36), 13223-13228. |
| Melaleuca quinquenervia | | Plantae | Sevillano Garcia Mayeya, L. (2010). The Effects of Biological Control Agents on Population Growth and Spread of Melaleuca quinquenervia. |
| Carlina vulgaris | | Plantae | Childs DZ, Rees M, Rose KE, Grubb PJ & Ellner SP (2003) Evolution of complex floweirng strategies: an age- and size-structured integral projection model. Proceedings: Biological Sciences 270: 1829-1838 |

| | | |
|---|---|---|
| Carlina vulgaris | Plantae | Childs DZ, Rees M, Rose KE, Grubb PY & Ellner SP (2004) Evolution of size-dependent flowering in a variable environment: construction and analysis of a stochastic integral projection model. Proceedings: Biological Sciences 271: 425-434 |
| Carlina vulgaris | Plantae | Rees M, Childs DZ, Metcalf JC, Rose KE, Sheppard AW & Grubb PJ (2006) Seed dormancy and delayed flowering in monocarpic plants: selective interactions in a stochastic environment. The American Naturalist 168: 53-71 |
| Carduus nutans | Plantae | Rees M, Childs DZ, Metcalf JC, Rose KE, Sheppard AW & Grubb PJ (2006) Seed dormancy and delayed flowering in monocarpic plants: selective interactions in a stochastic environment. The American Naturalist 168: 53-71 |
| Marmota ?aviventris | Animalia | Rees, M., Childs, D. Z., & Ellner, S. P. (2014). Building integral projection models: a user's guide. Journal of Animal Ecology, 83(3), 528-545. |
| Ovis aries | Animalia | Rees, M., Childs, D. Z., & Ellner, S. P. (2014). Building integral projection models: a user's guide. Journal of Animal Ecology, 83(3), 528-545. |
| Mammillaria dixanthocentron | Plantae | González EJ & Martorell C (2013) Reconstructing shifts in vital rates driven by long-term environmental change: a new demographic method based on readily available data. Ecology and Evolution 3: 2273–2284 |
| Cryptomeria japonica | Plantae | Matsushita, M., Takata, K., Hitsuma, G., Yagihashi, T., Noguchi, M., Shibata, M., & Masaki, T. (2015). A novel growth model evaluating age–size effect on long?term trends in tree growth. Functional Ecology. |
| Mammillaria gaumeri | Plantae | Ferrer-Cervantes, M.E., Méndez-González, M.E., Quintana-Ascencio, P.-F., Dorantes, A., Dzib, G. & Durán, R. (2012) Population dynamics of the cactus Mammillaria gaumeri: an integral projection model approach. Population Ecology, 54, 321-334. |
| Oenothera glazioviana | Plantae | Rees, M., & Rose, K. E. (2002). Evolution of flowering strategies in Oenothera glazioviana: an integral projection model approach. Proceedings of the Royal Society of London B: Biological Sciences, 269(1499), 1509-1515. |
| Prenanthes roanensis | Plantae | Aikens, M. L. (2013). Population Dynamics Across the Range of the Southern Appalachian Endemic Plant, Prenanthes Roanensis (Doctoral dissertation, University of Virginia). |
| Monstastraea annularis | Animalia | Burgess, H. R. (2011). Integral Projection Models and analysis of patch dynamics of the reef building coral Monstastraea annularis. |
| Sistrurus catenatus | Animalia | Jones, P. C. (2012). Demographic analysis and integral projection modeling of the Eastern Massasauga (Sistrurus catenatus catenatus) (Doctoral dissertation, NORTHERN ILLINOIS UNIVERSITY). |
| Marmota flaviventris | Animalia | Ozgul A, Childs DZ, Oli MK, Armitage KB, Blumstein DT, Olson LE, Tuljapurkar S & Coulson T (2010) Coupled dynamics of body mass and population growth in response to environmental change. Nature 466: 482-485 |
| Pedicularis lanceolata | Plantae | Record, S. (2010). Conservation while under invasion: Insights from a rare hemiparasitic plant, swamp lousewort (Pedicularis lanceolata Michx.). |
| Cirsium canescens | Plantae | Briggs J, Dabbs K, Holm M, Lubben J, Rebarber R, Tenhumberg B & Riser-Espinoza D (2010) Structured population dynamics: an introdution to integral modeling. Mathematical Magazine 83: 243-257 |
| Primula farinosa | Plantae | von Euler T, Ågren J & Ehrlén J (2013) Environmental context influences both the intensity of seed predation and plant demographic sensitivity to attack. Ecology in press |
| Annamocarya sinensis | Plantae | Zuidema PA, Jongejans E, Chien PD, During HJ & Schieving F (2010) Integral projection models for trees: a new parameterization method and a validation of model output. Journal of Ecology 98: 345-355 |
| Calocedrus macrolepis | Plantae | Zuidema PA, Jongejans E, Chien PD, During HJ & Schieving F (2010) Integral projection models for trees: a new parameterization method and a validation of model output. Journal of Ecology 98: 345-355 |

| | | | |
|---|---|---|---|
| Dacrydium elatum | | Plantae | Zuidema PA, Jongejans E, Chien PD, During HJ & Schieving F (2010) Integral projection models for trees: a new parameterization method and a validation of model output. Journal of Ecology 98: 345-355 |
| Manglietia fordiana | | Plantae | Zuidema PA, Jongejans E, Chien PD, During HJ & Schieving F (2010) Integral projection models for trees: a new parameterization method and a validation of model output. Journal of Ecology 98: 345-355 |
| Parashorea chinensis | | Plantae | Zuidema PA, Jongejans E, Chien PD, During HJ & Schieving F (2010) Integral projection models for trees: a new parameterization method and a validation of model output. Journal of Ecology 98: 345-355 |
| Pinus kwangtungensis | | Plantae | Zuidema PA, Jongejans E, Chien PD, During HJ & Schieving F (2010) Integral projection models for trees: a new parameterization method and a validation of model output. Journal of Ecology 98: 345-355 |
| Campanula thyrsoides | | Plantae | Kuss P, Rees M, Ægisdóttir HH, Ellner SP & Stöcklin J (2008) Evolutionary demography of long-lived monocarpic perennials: a time-lagged integral projection model. Journal of Ecology 96: 821-832 |
| Coryphantha robbinsorum | | Plantae | de Valpine P (2009) Stochastic development in biologically structured population models. Ecology 90: 2889-2901 |
| Artemisia tripartita | | Plantae | Alder PB, Dalgleish HJ & Ellner SP (2012) Forecasting plant community impacts of climate variability and change: when do competitive interactions matter? Journal of Ecology 100: 478-487 |
| Pseudoroegneria spicata | | Plantae | Alder PB, Dalgleish HJ & Ellner SP (2012) Forecasting plant community impacts of climate variability and change: when do competitive interactions matter? Journal of Ecology 100: 478-487 |
| Hesperostipa comata | | Plantae | Alder PB, Dalgleish HJ & Ellner SP (2012) Forecasting plant community impacts of climate variability and change: when do competitive interactions matter? Journal of Ecology 100: 478-487 |
| Poa secunda | | Plantae | Alder PB, Dalgleish HJ & Ellner SP (2012) Forecasting plant community impacts of climate variability and change: when do competitive interactions matter? Journal of Ecology 100: 478-487 |
| Artemisia tripartita | | Plantae | Adler PB, Ellner SP & Levine JM (2010) Coexistence of perennial plants: an embarrassment of niches. Ecology Letters 13: 1019-1029 |
| Pseudoroegneria spicata | | Plantae | Adler PB, Ellner SP & Levine JM (2010) Coexistence of perennial plants: an embarrassment of niches. Ecology Letters 13: 1019-1029 |
| Hesperostipa comata | | Plantae | Adler PB, Ellner SP & Levine JM (2010) Coexistence of perennial plants: an embarrassment of niches. Ecology Letters 13: 1019-1029 |
| Poa secunda | | Plantae | Adler PB, Ellner SP & Levine JM (2010) Coexistence of perennial plants: an embarrassment of niches. Ecology Letters 13: 1019-1029 |
| Acropora hyacinthus | | Animalia | Edmunds, P. J., Burgess, S. C., Putnam, H. M., Baskett, M. L., Bramanti, L., Fabina, N. S., ... & Gates, R. D. (2014). Evaluating the causal basis of ecological success within the scleractinia: an integral projection model approach. Marine Biology, 161(12), 2719-2734. |
| Chamaedorea elegans | | Plantae | Jansen M, Zuidema PA, Anten NPR & Martinez-Ramos M (2012) Strong persistent growth differences govern individual performance and populaton dynamics in a tropical forest understory palm. Journal of Ecoloy 100: 1224-1232 |
| Poecilia reticulata | | Animalia | Basar RD, Lopez-Sepulcre A, Reznick DN & Travis J (2013) Experimental evidence for density-dependent regulation and selection on Trinidadian guppy life history. American Naturalist 181: 25-38 |
| Poecilia reticulata | | Animalia | Bassar, R. D., Heatherly, T., Marshall, M. C., Thomas, S. A., Flecker, A. S., & Reznick, D. N. (2015). Population size?structure?dependent fitness and ecosystem consequences in Trinidadian guppies. Journal of Animal Ecology. |

| | | |
|---|---|---|
| Poecilia reticulata | Animalia | Bassar, R. D., Lopez-Sepulcre, A., Reznick, D. N., & Travis, J. (2013). Experimental evidence for density-dependent regulation and selection on Trinidadian guppy life histories. The American Naturalist, 181(1), 25-38. |
| Cirsium canescens | Plantae | Rebarber R, Tenhumberg B & Townley S (2012) Global asymptotic stability of density dependent integral population projection models. Theoretical Population Biology 81: 81-87 |
| Cryptantha flava | Plantae | Salguero-Gómez R, Siewert W, Casper BB & Tielbörger K (2012) A demographic approach to study effects of climate change in desert plants. Philosophical Transaction of the Royal Society Series B 367: 3100-3114 |
| Cryptantha flava | Plantae | NA |
| Succisa pratensis | Plantae | van der Meer S, Dahlgren JP, Mildén M & Ehrlén J (2013) Differential effects of abandonment on the demography of the grassland perennial Succisa pratensis. Population Ecology, in press |
| Cirsium palustre | Plantae | Ramula S, Rees M & Buckley YM (2009) Integral projection models perform beter for small demographic data sets than matrix populatio models: a case study of two perennial herbs. Journal of Applied Ecology 46: 1048-1053 |
| Primula veris | Plantae | Ramula S, Rees M & Buckley YM (2009) Integral projection models perform beter for small demographic data sets than matrix populatio models: a case study of two perennial herbs. Journal of Applied Ecology 46: 1048-1053 |
| Cirsium canescens | Plantae | Eager, E. A. (2012). Modeling and mathematical analysis of plant models in ecology (Doctoral dissertation, University of Nebraska). |
| Helianthus annuus | Plantae | Eager, E. A., Rebarber, R., & Tenhumberg, B. (2014). Modeling and Analysis of a Density-Dependent Stochastic Integral Projection Model for a Disturbance Specialist Plant and Its Seed Bank. Bulletin of mathematical biology, 76(7), 1809-1834. |
| Cirsium palustre | Plantae | Eager, E. A., Rebarber, R., & Tenhumberg, B. (2014). Global asymptotic stability of plant-seed bank models. Journal of mathematical biology, 69(1), 1-37. |
| Hedysarum laeve | Plantae | Li, S. L., Yu, F. H., Werger, M. J., Dong, M., During, H. J., & Zuidema, P. A. (2015). Mobile dune fixation by a fast-growing clonal plant: a full life-cycle analysis. Scientific reports, 5. |
| Carlina vulgaris | Plantae | Rees M & Ellner SP (2009) Integral projection models for populations in temporally varying environments. Ecological Monographs 79: 575-594 |
| Lupinus polyphyllus | Plantae | Ramula, S. (2014). Linking vital rates to invasiveness of a perennial herb. Oecologia, 174(4), 1255-1264. |
| Vaccinium myrtillus | Plantae | Hegland SJ, Jongejans E & Rydgren K (2010) Investigating the interaction between ungulate grazing and resource effects on Vaccinium myrtillus populations with integral projection models. Oecology 163: 695-706 |
| Urocitellus columbianus | Animalia | Schindler S, Neuhaus P, Gaillard J-M & Coulson T (2013) The influence of nonrandom mating on population growth. American Naturalist 182: 28–41 |
| Ovis aries | Animalia | Coulson T (2012) Integral projection models, their construction and use in posing hypotheses in ecology. Oikos 121: 1337-1350 |
| Canis lupus | Animalia | Coulson T (2001) Modeling effects of environmental change on wolf population dynamics, trait evolution, and life history. Science 334: 1275-1278 |
| Ovis aries | Animalia | Coulson T, Tuljapurkar S & Childs DZ (2010) Using evolutionary demography to link life history theory, quantitative genetics and population ecology. Journal of Animal Ecology 79: 1226-1240 |
| Ovis aries | Animalia | Coulson, T., Tuljapurkar, S., & Childs, D. Z. (2010). Using evolutionary demography to link life history theory, quantitative genetics and population ecology. Journal of Animal Ecology, 79(6), 1226-1240. |
| Rhinella marina | Animalia | Perkins TA, Phillips BL, Baskett ML & Hastings A (2013) Evolution of dispersal and life history interact to drive accelerating spread of an invasive species. Ecology Letters 16: 1079–1087 |

| | | |
|---|---|---|
| Opuntia imbricata | Plantae | Miller TEX, Louda SM, Rose KA & Eckberg JO (2009) Impacts of insect herbivory on cactus population dynamics: experimental demography across an environmental gradient. Ecological Monographs 79: 155-172 |
| Orchis purpurea | Plantae | Miller TEX, Williams JL, Jongejans E, Brys R & Jacquemyn H (2012) Evolutionary demography of iteroparous plants: incorporating non-lethal costs of reproduction into integral projection models. Proc Roy Soc B, 279: 2831-2840 |
| Miliusa horsfieldii | Plantae | Caughlin TT, Ferguson JM, Lichstein JW, Zuidema PA, Bunyavejchewin S, Levey DJ. 2015 Loss of animal seed dispersal increases extinction risk in a tropical tree species due to pervasive negative density dependence across life stages. Proc. R. Soc. B 282: 20142095. http://dx.doi.org/10.1098/rspb.2014.2095 |
| Esox lucius | Animalia | Vindenes, Y., Edeline, E., Ohlberger, J., Langangen, Ø., Winfield, I. J., Stenseth, N. C., & Vøllestad, L. A. (2014). Effects of climate change on trait-based dynamics of a top predator in freshwater ecosystems. The American Naturalist, 183(2), 243-256. |
| Berberis thunbergii | Plantae | Merow, C., Bois, S. T., Allen, J. M., Xie, Y., Silander Jr., J. A. (2017). Climate change both facilitates and inhibits invasive plant ranges in New England. PNAS Vol. 114, Issue 16. |
| Cirsium altissimum | Plantae | Tenhumberg, B., Suwa, T., Tyre, A. J., Russell, F. L., & Louda, S. M. (2015). Integral projection models show exotic thistle is more limited than native thistle by ambient competition and herbivory. Ecosphere, 6(4), art69. |
| NDY | Animalia | Jason Matthiopoulos, John Fieberg, Geert Aarts, Hawthorne L. Beyer, Juan M. Morales, and Daniel T. Haydon 2015. Establishing the link between habitat selection and animal population dynamics. Ecological Monographs 85:413–436. http://dx.doi.org/10.1890/14-2244.1 |
| Bouteloua eriopoda | Plantae | Chengjin Chu and Peter B. Adler 2015. Large niche differences emerge at the recruitment stage to stabilize grassland coexistence. Ecological Monographs 85:373–392. http://dx.doi.org/10.1890/14-1741.1 |
| Bouteloua rothrockii | Plantae | Chengjin Chu and Peter B. Adler 2015. Large niche differences emerge at the recruitment stage to stabilize grassland coexistence. Ecological Monographs 85:373–392. http://dx.doi.org/10.1890/14-1741.1 |
| Artemisia tripartita | Plantae | Chengjin Chu and Peter B. Adler 2015. Large niche differences emerge at the recruitment stage to stabilize grassland coexistence. Ecological Monographs 85:373–392. http://dx.doi.org/10.1890/14-1741.1 |
| Pseudoroegneria spicata | Plantae | Chengjin Chu and Peter B. Adler 2015. Large niche differences emerge at the recruitment stage to stabilize grassland coexistence. Ecological Monographs 85:373–392. http://dx.doi.org/10.1890/14-1741.1 |
| Hesperostipa comata | Plantae | Chengjin Chu and Peter B. Adler 2015. Large niche differences emerge at the recruitment stage to stabilize grassland coexistence. Ecological Monographs 85:373–392. http://dx.doi.org/10.1890/14-1741.1 |
| Poa secunda | Plantae | Chengjin Chu and Peter B. Adler 2015. Large niche differences emerge at the recruitment stage to stabilize grassland coexistence. Ecological Monographs 85:373–392. http://dx.doi.org/10.1890/14-1741.1 |
| Bouteloua curtipendula | Plantae | Chengjin Chu and Peter B. Adler 2015. Large niche differences emerge at the recruitment stage to stabilize grassland coexistence. Ecological Monographs 85:373–392. http://dx.doi.org/10.1890/14-1741.1 |
| Bouteloua hirsuta | Plantae | Chengjin Chu and Peter B. Adler 2015. Large niche differences emerge at the recruitment stage to stabilize grassland coexistence. Ecological Monographs 85:373–392. http://dx.doi.org/10.1890/14-1741.1 |
| Schizachyrium scoparium | Plantae | Chengjin Chu and Peter B. Adler 2015. Large niche differences emerge at the recruitment stage to stabilize grassland coexistence. Ecological Monographs 85:373–392. http://dx.doi.org/10.1890/14-1741.1 |

| | | |
|---|---|---|
| Bouteloua gracilis | Plantae | Chengjin Chu and Peter B. Adler 2015. Large niche differences emerge at the recruitment stage to stabilize grassland coexistence. Ecological Monographs 85:373–392. http://dx.doi.org/10.1890/14-1741.1 |
| Pascopyrum smithii | Plantae | Chengjin Chu and Peter B. Adler 2015. Large niche differences emerge at the recruitment stage to stabilize grassland coexistence. Ecological Monographs 85:373–392. http://dx.doi.org/10.1890/14-1741.1 |
| Sporobolus flexuosus | Plantae | Chengjin Chu and Peter B. Adler 2015. Large niche differences emerge at the recruitment stage to stabilize grassland coexistence. Ecological Monographs 85:373–392. http://dx.doi.org/10.1890/14-1741.1 |
| Ferocactus wislizeni | Plantae | Ford, K. R., Ness, J. H., Bronstein, J. L., & Morris, W. F. (2015). The demographic consequences of mutualism: ants increase host-plant fruit production but not population growth. Oecologia, 1-12. |
| Festuca arizonica | Plantae | NA |
| Poecilia reticulata | Animalia | Bassar, R. D., Childs, D. Z., Rees, M., Tuljapurkar, S., Reznick, D. N., Coulson, T. (2016). The effects of asymmetric competition on the life history of Trinidadian guppies. Ecology Letters. |
| Ovis canadensis | Animalia | Pigeon, G., Festa-Bianchet, M., Coltman, D. W., Pelletier, F. (2016). Intense selective hunting leads to artificial evolution in horn size. Evolutionary Applications. |
| Salvelinus fontinalis | Animalia | Bassar, R. D., Letcher, B. H., Nislow, K. H., Whiteley, A. R. (2016). Changes in seasonal climate outpace compensatory density-dependence in eastern brook trout. Global Change Biology. |
| Achyranthes japonica | Plantae | Schwartz, L. M., Gibson, D. J., Young, B. G. (2016). Using integral projection models to compare population dynamics of four closely related species. Population Ecology. |
| Amaranthus palmeri | Plantae | Schwartz, L. M., Gibson, D. J., Young, B. G. (2016). Using integral projection models to compare population dynamics of four closely related species. Population Ecology. |
| Amaranthus tuberculatus | Plantae | Schwartz, L. M., Gibson, D. J., Young, B. G. (2016). Using integral projection models to compare population dynamics of four closely related species. Population Ecology. |
| Iresine rhizomatosa | Plantae | Schwartz, L. M., Gibson, D. J., Young, B. G. (2016). Using integral projection models to compare population dynamics of four closely related species. Population Ecology. |
| Aquila fasciata | Animalia | Lieury, N., Besnard, A., Ponchon, C., Ravayrol, A., Millon, A. (2016). Geographically isolated but demographically connected: Immigration supports efficient conservation actions in the recovery of a range-margin population of the Bonelli's eagle in France. Biological Conservation 195. |
| Cryptantha flava | Plantae | Schreiber, S., & Ross, N. (2016). Individual-based Integral Projection Models: The role of size-structure on extinction risk and establishment success. Methods in Ecology and Evolution. |
| Veronica officinalis | Plantae | Olsen, S. L., Töpper, J. P., Skarpaas, O., Vandvik, V., Klanderud, K. (2016). From facilitation to competition: temperature-driven shift in dominant plant interactions affects population dynamics in semi-natural grasslands. Global Change Biology. |
| Veronica alpina | Plantae | Olsen, S. L., Töpper, J. P., Skarpaas, O., Vandvik, V., Klanderud, K. (2016). From facilitation to competition: temperature-driven shift in dominant plant interactions affects population dynamics in semi-natural grasslands. Global Change Biology. |
| Viola palustris | Plantae | Olsen, S. L., Töpper, J. P., Skarpaas, O., Vandvik, V., Klanderud, K. (2016). From facilitation to competition: temperature-driven shift in dominant plant interactions affects population dynamics in semi-natural grasslands. Global Change Biology. |

| | | | |
|---|---|---|---|
| Viola biflora | | Plantae | Olsen, S. L., Töpper, J. P., Skarpaas, O., Vandvik, V., Klanderud, K. (2016). From facilitation to competition: temperature-driven shift in dominant plant interactions affects population dynamics in semi-natural grasslands. Global Change Biology. |
| Cryptantha flava | | Plantae | González, E. J., Martorell, C., & Bolker, B. M. (2016). Inverse estimation of integral projection model parameters using time series of population?level data.Methods in Ecology and Evolution, 7(2), 147-156. |
| Parus major | | Animalia | Childs, D. Z., Sheldon, B. C., & Rees, M. (2016). The evolution of labile traits in sex?and age?structured populations. Journal of Animal Ecology, 85(2), 329-342. |
| Fraxinus excelsior | | Plantae | Needham, J., Merow, C., Butt, N., Malhi, Y., Marthews, T. R., Morecroft, M., & McMahon, S. M. (2016). Forest community response to invasive pathogens: the case of ash dieback in a British woodland. Journal of Ecology, 104(2), 315-330. |
| Prenanthes_roanensis | | Plantae | Aikens, M. L., & Roach, D. A. (2014). Population dynamics in central and edge populations of a narrowly endemic plant. Ecology, 95(7), 1850-1860. |
| Himantoglossum_hircinum | | Plantae | Van der Meer, S., Jacquemyn, H., Carey, P. D., Jongejans, E. (2016). Recent range expansion of a terrestrial orchid corresponds with climate-driven variation in ist population dynamics. Oecologia. |
| Grus americana | | Animalia | Wilson, S., Gil-Weir, K. C., Clark, R. G., Robertson, G. J., & Bidwell, M. T. (2016). Integrated population modeling to assess demographic variation and contributions to population growth for endangered whooping cranes. Biological Conservation, 197, 1-7. |
| Opuntia imbricata | | Plantae | Elderd, B. D., & Miller, T. E. (2015). Quantifying demographic uncertainty: Bayesian methods for Integral Projection Models (IPMs). Ecological Monographs. |
| Oenocarpus bataua | | Plantae | Isaza, C., Martorell, C., Cevallos, D., Galeano, G., Valencia, R., Balslev, H. (2016). Demography of Oenocarpus bataua and implications for sustainable harvest of its fruit in western Amazon. Population Ecology. |
| Cotyledon orbiculata | | Plantae | Pannell, J. L. (2016). Climatic limitation of alien weeds in New Zealand: enhancing species distribution models with field data (Doctoral dissertation, Lincoln University). |
| Geum radiatum | | Plantae | Ulrey, C., Quintana-Ascencio, P., Kauffman, G., Smith, A. B., Menges, E. S. (2016). Life at the top: Long-term demography, microclimatic refugia, and responses to climate change for a high-elevation southern Appalachian endemic plant. Biological Conservation 200 (2016) 80-92. |
| Ivesia lycopodioides var. scandularis | | Plantae | Oldfather, M. F., & Ackerly, D. D. (2018). Microclimate and demography interact to shape stable population dynamics across the range of an alpine plant. New Phytologist. |
| Fumana procumbens | | Plantae | Dahlgren, J. P., Bengtsson, K., Ehrlen J. (2016). The demography of climate-driven and density- regulated population dynamics in a perennial plant. Ecology, 97(4), 899-907 |
| Esox lucius | | Animalia | Vindenes, Y., Langangen, Ø., Winfield, I. J., Vøllestad, L. A. (2016). Fitness consequences of early life conditions and maternal size effects in a freshwater top predator. Journal of Animal Ecology, (85), 692-704 |
| Crassostrea gigas | | Animalia | Moore, J. L., Lipcius, R. N., Puckett, B., Schreiber, S. J. (2016). The demographic consequences of growing older and bigger oyster populations. Ecological Applications. |
| Callitris intratropica | | Plantae | Trauernicht, C., Murphy, B. P., Prior, L. D., Lawes, M. J., & Bowman, D. M. (2016). Human-Imposed, Fine-Grained Patch Burning Explains the Population Stability of a Fire-Sensitive Conifer in a Frequently Burnt Northern Australia Savanna. Ecosystems, 1-14. |
| Rana mucosa | | Animalia | Wilber, M. Q., Langwig, K. E., Kilpatrick, A. M., McCallum, H. I., & Briggs, C. J. (2016). Integral Projection Models for host–parasite systems with an application to amphibian chytrid fungus. Methods in Ecology and Evolution. |

| | | |
|---|---|---|
| Cervus elaphus | Animalia | Pozo, R. A., Schindler, S., Cubaynes, S., Cusack, J. J., Coulson, T., Malo, A. F., Modeling the Impact of Selective Harvesting on Red Deer Antlers. J Wildlife Manage. |
| Lindera_benzoin | Plantae | Merow, C., Bois, S. T., Allen, J. M., Xie, Y., Silander Jr., J. A. (2017). Climate change both facilitates and inhibits invasive plant ranges in New England. PNAS Vol. 114, Issue 16. |
| Dacrydium elatum | Plantae | Snyder, E. S., Ellner, S. P. (2016). We Happy Few: Using Structured Population Models to Identidy the Decisive Events in the Lives of Exceptional Individuals. The American Naturalist, Vol. 188, No. 2. |
| Liatris ohlingerae | Plantae | Tye, M. R., Menges, E. S., Weekley, C. (2016). A demographic ménage à trois: interactions between disturbances both amplify and dampen populatin dynamics if an endemic plant. Journal of Ecology. |
| Sebastes mystinus | Animalia | White, J. W., Nickols, K. J., Malone, D., Carr, M. H., Starr, R. M., Cordoleani, F., Baskett, M. L., Hastings, A., Botsford, L. W. (2016). Fitting state-space integral projection models to size-structured time series data to estimate unknown parameters. Ecological Applications. |
| Sebastes carnatus | Animalia | White, J. W., Nickols, K. J., Malone, D., Carr, M. H., Starr, R. M., Cordoleani, F., Baskett, M. L., Hastings, A., Botsford, L. W. (2016). Fitting state-space integral projection models to size-structured time series data to estimate unknown parameters. Ecological Applications. |
| Opuntia imbricata | Plantae | Compagnoni, A., Bibian, A. J., Ochocki, B. M., Rogers, H. S., Schultz, E. L., Sneck, M. E., Elderd, B. D., Iler, A. M., Inouye, D. W., Jacquemyn, H., Miller, T. E. X. (2016). The effect of demographic correlations on the stochastic population dynamics of perennial plants. Ecological Monographs. |
| Orchis purpurea | Plantae | Compagnoni, A., Bibian, A. J., Ochocki, B. M., Rogers, H. S., Schultz, E. L., Sneck, M. E., Elderd, B. D., Iler, A. M., Inouye, D. W., Jacquemyn, H., Miller, T. E. X. (2016). The effect of demographic correlations on the stochastic population dynamics of perennial plants. Ecological Monographs. |
| Pinus halepensis | Plantae | García-Callejas, D., Molowny-Horas, R., Retana, J. (2016). Projecting the distribution and abundance of Mediterranean tree species under climate change: a demographic approach. Journal of Plant Ecology. |
| Artemisia tripartita | Plantae | Ellner, S. P., Snyder, R. E., Adler, P. B. (2016). How to quantify the temporal storage effect using simulations instead of math. Ecology Letters. |
| Hesperostipa comata | Plantae | Ellner, S. P., Snyder, R. E., Adler, P. B. (2016). How to quantify the temporal storage effect using simulations instead of math. Ecology Letters. |
| Poa secunda | Plantae | Ellner, S. P., Snyder, R. E., Adler, P. B. (2016). How to quantify the temporal storage effect using simulations instead of math. Ecology Letters. |
| Pseudoroegneria spicata | Plantae | Ellner, S. P., Snyder, R. E., Adler, P. B. (2016). How to quantify the temporal storage effect using simulations instead of math. Ecology Letters. |
| Cirsium canescens | Plantae | Eager, E. A., Rebarber, R. (2016). Sensitivity and elasticity analysis of a Lur'e system used to model a population subject to density-dependent reproduction. Mathematical Biosciences. |
| Zootoca vivipara | Animalia | Jaffré, M.; Le Galliard, J.F. (2016). Population viability analysis of plant and animal populations with stochastic integral projection models. Oecologia |
| Rhizoglyphus robini | Animalia | Smallegange, I. M., Caswell, H., Toorians, M. E. M., de Roos, A. M. (2016). Mechanistic description of population dynamics using dynamic energy budget theory incorporated into integral projection models. Methods in Ecology and Evolution. |
| Manta alfredi | Animalia | Smallegange, I. M., Caswell, H., Toorians, M. E. M., de Roos, A. M. (2016). Mechanistic description of population dynamics using dynamic energy budget theory incorporated into integral projection models. Methods in Ecology and Evolution. |

| | | |
|---|---|---|
| Drosophyllum lusitanicum | Plantae | Paniw, M., Quintana-Ascencio, P. F., Ojeda, F., Salguero-Gómez, R. (2016). Accounting for uncertainty in dormant life stages in stochastic demographic models. Oikos. |
| Bouteloua gracilis | Plantae | Tredennick, A. T., Hooten, M. B., Adler, P. B. (2016). Do we need demographic data to forecast plant population dynamics? bioRxiv. |
| Hesperostipa comata | Plantae | Tredennick, A. T., Hooten, M. B., Adler, P. B. (2016). Do we need demographic data to forecast plant population dynamics? bioRxiv. |
| Pascopyrum smithii | Plantae | Tredennick, A. T., Hooten, M. B., Adler, P. B. (2016). Do we need demographic data to forecast plant population dynamics? bioRxiv. |
| Poa secunda | Plantae | Tredennick, A. T., Hooten, M. B., Adler, P. B. (2016). Do we need demographic data to forecast plant population dynamics? bioRxiv. |
| Tillandsia macdougallii | Plantae | Ticktin, T., Mondragón, D., and Gaoue, O.G. (2016). Host genus and rainfall drive the population dynamics of a vascular epiphyte. Ecosphere 7.11 |
| Pometia spp. | Plantae | Murdjoko, A., Marsono, D., Sandono, R., Hadisusanto, S. (2016). Population Dynamics of Pometia for The Period of Post-Selective Logging in Tropical Rainforest, Southern Papua, Indonesia. Biosaintifika: Journal of Biology & Biology Education 8 (3). 320-329. |
| Cyprinodontiformes_sp. | Animalia | Bassar, R. D., Simon, T., Roberts, W., Travis, J., Reznick, D. N. (2016). The evolution of coexistence: Reciprocal adaptation promotes the assembly of a simple community. Evolution. |
| Poecilia reticulata | Animalia | Bassar, R. D., Simon, T., Roberts, W., Travis, J., Reznick, D. N. (2016). The evolution of coexistence: Reciprocal adaptation promotes the assembly of a simple community. Evolution. |
| Pinus halepensis | Plantae | Molowny Horas, R., & Espelta, J.M. (2013). Modelos integrales de proyección como instrumentos para la gestón medioambiental forestal. En J.A. Blanco (Ed.). Aplicaciones de modelos ecológicos a la gesti ón de recursos naturales . (pp. 125-140). Barcelona: OmniaScience. |
| Drosophyllum lusitanicum | Plantae | Paniw, M., Quintana-Ascencio, P. F., Ojeda, F., Salguero-Gómez, R. (2017). Interacting livestock and fire may both threaten and increase viability of a fire-adapted Mediterranean carnivorous plant. Journal of Applied Ecology. |
| Alchornea costaricensis | Plantae | Bruijning, M., Visser, M. D., Muller-Landau, H. C., Wright, S. J., Comita, L. S., Hubbell, S. P., De Kroon, H., Jongejans, E. (2017). Surviving in a Cosexual World: A Cost-Benefit Analysis of Dioecy in Tropical Trees. The American Naturalist. Vol. 189, No. 3. |
| Cecropia insignis | Plantae | Bruijning, M., Visser, M. D., Muller-Landau, H. C., Wright, S. J., Comita, L. S., Hubbell, S. P., De Kroon, H., Jongejans, E. (2017). Surviving in a Cosexual World: A Cost-Benefit Analysis of Dioecy in Tropical Trees. The American Naturalist. Vol. 189, No. 3. |
| Cecropia obtusifolia | Plantae | Bruijning, M., Visser, M. D., Muller-Landau, H. C., Wright, S. J., Comita, L. S., Hubbell, S. P., De Kroon, H., Jongejans, E. (2017). Surviving in a Cosexual World: A Cost-Benefit Analysis of Dioecy in Tropical Trees. The American Naturalist. Vol. 189, No. 3. |
| Simarouba amara | Plantae | Bruijning, M., Visser, M. D., Muller-Landau, H. C., Wright, S. J., Comita, L. S., Hubbell, S. P., De Kroon, H., Jongejans, E. (2017). Surviving in a Cosexual World: A Cost-Benefit Analysis of Dioecy in Tropical Trees. The American Naturalist. Vol. 189, No. 3. |
| Pouteria reticulata | Plantae | Bruijning, M., Visser, M. D., Muller-Landau, H. C., Wright, S. J., Comita, L. S., Hubbell, S. P., De Kroon, H., Jongejans, E. (2017). Surviving in a Cosexual World: A Cost-Benefit Analysis of Dioecy in Tropical Trees. The American Naturalist. Vol. 189, No. 3. |

| | | |
|---|---|---|
| Protium tenuifolium | Plantae | Bruijning, M., Visser, M. D., Muller-Landau, H. C., Wright, S. J., Comita, L. S., Hubbell, S. P., De Kroon, H., Jongejans, E. (2017). Surviving in a Cosexual World: A Cost-Benefit Analysis of Dioecy in Tropical Trees. The American Naturalist. Vol. 189, No. 3. |
| Triplaris cumingiana | Plantae | Bruijning, M., Visser, M. D., Muller-Landau, H. C., Wright, S. J., Comita, L. S., Hubbell, S. P., De Kroon, H., Jongejans, E. (2017). Surviving in a Cosexual World: A Cost-Benefit Analysis of Dioecy in Tropical Trees. The American Naturalist. Vol. 189, No. 3. |
| Virola sebifera | Plantae | Bruijning, M., Visser, M. D., Muller-Landau, H. C., Wright, S. J., Comita, L. S., Hubbell, S. P., De Kroon, H., Jongejans, E. (2017). Surviving in a Cosexual World: A Cost-Benefit Analysis of Dioecy in Tropical Trees. The American Naturalist. Vol. 189, No. 3. |
| Bouteloua eriopoda | Plantae | Tredennick, A. T., De Mazancourt, C., Loreau, M., Adler, P. B. (2017). Environmental responses, not species interactions, determine synchrony of dominant species in semiarid grasslands. Ecology. |
| Sporobolus ?exuosus | Plantae | Tredennick, A. T., De Mazancourt, C., Loreau, M., Adler, P. B. (2017). Environmental responses, not species interactions, determine synchrony of dominant species in semiarid grasslands. Ecology. |
| Bouteloua eriopoda | Plantae | Tredennick, A. T., De Mazancourt, C., Loreau, M., Adler, P. B. (2017). Environmental responses, not species interactions, determine synchrony of dominant species in semiarid grasslands. Ecology. |
| Bouteloua rothrockii | Plantae | Tredennick, A. T., De Mazancourt, C., Loreau, M., Adler, P. B. (2017). Environmental responses, not species interactions, determine synchrony of dominant species in semiarid grasslands. Ecology. |
| Bouteloua curtipendula | Plantae | Tredennick, A. T., De Mazancourt, C., Loreau, M., Adler, P. B. (2017). Environmental responses, not species interactions, determine synchrony of dominant species in semiarid grasslands. Ecology. |
| Bouteloua hirsuta | Plantae | Tredennick, A. T., De Mazancourt, C., Loreau, M., Adler, P. B. (2017). Environmental responses, not species interactions, determine synchrony of dominant species in semiarid grasslands. Ecology. |
| Schizachyrium scoparium | Plantae | Tredennick, A. T., De Mazancourt, C., Loreau, M., Adler, P. B. (2017). Environmental responses, not species interactions, determine synchrony of dominant species in semiarid grasslands. Ecology. |
| Bouteloua gracilis | Plantae | Tredennick, A. T., De Mazancourt, C., Loreau, M., Adler, P. B. (2017). Environmental responses, not species interactions, determine synchrony of dominant species in semiarid grasslands. Ecology. |
| Hesperostipa comata | Plantae | Tredennick, A. T., De Mazancourt, C., Loreau, M., Adler, P. B. (2017). Environmental responses, not species interactions, determine synchrony of dominant species in semiarid grasslands. Ecology. |
| Pascopyrum smithii | Plantae | Tredennick, A. T., De Mazancourt, C., Loreau, M., Adler, P. B. (2017). Environmental responses, not species interactions, determine synchrony of dominant species in semiarid grasslands. Ecology. |
| Poa secunda | Plantae | Tredennick, A. T., De Mazancourt, C., Loreau, M., Adler, P. B. (2017). Environmental responses, not species interactions, determine synchrony of dominant species in semiarid grasslands. Ecology. |
| Artemisia tripartita | Plantae | Tredennick, A. T., De Mazancourt, C., Loreau, M., Adler, P. B. (2017). Environmental responses, not species interactions, determine synchrony of dominant species in semiarid grasslands. Ecology. |
| Pseudoroegneria spicata | Plantae | Tredennick, A. T., De Mazancourt, C., Loreau, M., Adler, P. B. (2017). Environmental responses, not species interactions, determine synchrony of dominant species in semiarid grasslands. Ecology. |

| | | |
|---|---|---|
| Hesperostipa comata | Plantae | Tredennick, A. T., De Mazancourt, C., Loreau, M., Adler, P. B. (2017). Environmental responses, not species interactions, determine synchrony of dominant species in semiarid grasslands. Ecology. |
| Poa secunda | Plantae | Tredennick, A. T., De Mazancourt, C., Loreau, M., Adler, P. B. (2017). Environmental responses, not species interactions, determine synchrony of dominant species in semiarid grasslands. Ecology. |
| Gadus morhua | Animalia | Aalto, E. A., Baskett, M. L. (2017). Post-harvest recovery dynamics depend on predator specialization in size-selective fisheries. Marine Ecology Progress Series. Vol 564: 127-143. |
| Melanogrammus aeglefinus | Animalia | Aalto, E. A., Baskett, M. L. (2017). Post-harvest recovery dynamics depend on predator specialization in size-selective fisheries. Marine Ecology Progress Series. Vol 564: 127-143. |
| Merlangius merlangus | Animalia | Aalto, E. A., Baskett, M. L. (2017). Post-harvest recovery dynamics depend on predator specialization in size-selective fisheries. Marine Ecology Progress Series. Vol 564: 127-143. |
| Balanophyllia elegans | Animalia | Elahi, R., Sebens, K. P., De Leo, G. A. (2016). Ocean warming and the demography oi declines in coral body size. Marine Ecology Progress Series. Vol. 560: 147-158. |
| Calathea crotalifera | Plantae | Westerband, A. C., Horvitz, C. C. (2017). Photosynthetic rates influence the population dynamics of understory herbs in stochastic light environments. Ecology, 98 (2): 370-381. |
| Heliconia tortuosa | Plantae | Westerband, A. C., Horvitz, C. C. (2017). Photosynthetic rates influence the population dynamics of understory herbs in stochastic light environments. Ecology, 98 (2): 370-381. |
| Microtus ochrogaster | Animalia | Van Benthem, K. J., Froy, H., Coulson, T., Getz, L. L., Oli, M. K., Ozgul, A. (2017). Trait – demography relationships underlying small mammal population ?uctuations. Journal of Animal Ecology, 86: 348-358. |
| Mammillaria dixanthocentron | Plantae | Ureta, C., Martorell, C., Hortal, J., & Fornoni, J. (2012). Assessing extinction risks under the combined effects of climate change and human disturbance through the analysis of life-history plasticity. Perspectives in Plant Ecology, Evolution and Systematics, 14(6), 393-401. |
| Mammillaria hernandezii | Plantae | Ureta, C., Martorell, C., Hortal, J., & Fornoni, J. (2012). Assessing extinction risks under the combined effects of climate change and human disturbance through the analysis of life-history plasticity. Perspectives in Plant Ecology, Evolution and Systematics, 14(6), 393-401. |
| Astrocaryum chambira | Plantae | García, N., Zuidema, P. A., Galeano, G., & Bernal, R. (2016). Demography and sustainable management of two fiber?producing Astrocaryum palms in Colombia. Biotropica, 48(5), 598-607. |
| Astrocaryum standleyanum | Plantae | García, N., Zuidema, P. A., Galeano, G., & Bernal, R. (2016). Demography and sustainable management of two fiber?producing Astrocaryum palms in Colombia. Biotropica, 48(5), 598-607. |
| Ovis canadensis | Animalia | Manlove, Kezia, et al. "Disease introduction is associated with a phase transition in bighorn sheep demographics." Ecology 97.10 (2016): 2593-2602. |
| Nothofagus dombeyi | Plantae | Molowny?Horas, R., Suarez, M. L., & Lloret, F. (2017). Changes in the natural dynamics of Nothofagus dombeyi forests: population modeling with increasing drought frequencies. Ecosphere, 8(3), e01708. |
| Poa alsodes | Plantae | Chung, Y. A., Miller, T. E., & Rudgers, J. A. (2015). Fungal symbionts maintain a rare plant population but demographic advantage drives the dominance of a common host. Journal of Ecology. |
| Poa sylvestris | Plantae | Chung, Y. A., Miller, T. E., & Rudgers, J. A. (2015). Fungal symbionts maintain a rare plant population but demographic advantage drives the dominance of a common host. Journal of Ecology. |

| | | |
|---|---|---|
| Protea repens | Plantae | Merow, C., Latimer, A. M., Wilson, A. M., McMahon, S. M., Rebelo, A. G., Silander Jr, J. A. (2014). On using integral projecction modls to generate demographically driven predictions of species distributions: development and validation using sparse data. Ecography 32: 1167-1183 |
| Opuntia rastrera | Plantae | Ureta, C., Martorell, C., Cuervo-Robayo, A. P., Mandujano, M. C., Martínez-Meyer, E. (2018). Inferring space from time: On the relationship between demography and environmental suitability in the desert plan O. rastrera. Plos ONE 13(8): e0201543 |
| Ambrosia artemisiifolia | Plantae | Lommen, S. T. E., Jongejans, E., Leitsch-Vitalos, M., Tokarska-Guzik, B., Zalai, M., Müller-Schärer, H., Karrer, G. (2018). Time to cut: population models reveal how to mow invasive common ragweed cost-effectively. NeoBiota (39):: 53-78 |
| Ivesia lycopodioides A. Gray var. scandularis | Plantae | Oldfather, M. F. (2018). Population and Community Dynamics of Alpine Plants in a Changing Climate Across Topographically Heterogeneous Landscapes (Doctoral dissertation, UC Berkeley). |
| Turritis_glabra | Plantae | Merow, C., Bois, S. T., Allen, J. M., Xie, Y., Silander Jr., J. A. (2017). Climate change both facilitates and inhibits invasive plant ranges in New England. PNAS Vol. 114, Issue 16. |
| Daphnia magna | Animalia | Bruijning, M., ten Berge, A. C., & Jongejans, E. (2018). Population?level responses to temperature, density and clonal differences in Daphnia magna as revealed by integral projection modelling. Functional Ecology, 32(10), 2407-2422. |
| Pinus massoniana | Plantae | Yang, X., Li, S., Shen, B., Wu, Y., Sun, S., Liu, R., ... & Li, S. L. (2018). Demographic strategies of a dominant tree species in response to logging in a degraded subtropical forest in Southeast China. Annals of Forest Science, 75(3), 84. |
| Viola biflora L. var. rockiana | Plantae | Cui, H., Töpper, J. P., Yang, Y., Vandvik, V., & Wang, G. (2018). Plastic population effects and conservative leaf traits in a reciprocal transplant experiment simulating climate warming in the Himalayas. Frontiers in plant science, 9. |
| Acropora sp.; Pocillopora sp; Porites sp; | Animalia | Kayal, M., Lenihan, H. S., Brooks, A. J., Holbrook, S. J., Schmitt, R. J., & Kendall, B. E. (2018). Predicting coral community recovery using multi?species population dynamics models. Ecology letters. |
| Marmota flaviventer | Animalia | Maldonado?Chaparro, A. A., Blumstein, D. T., Armitage, K. B., & Childs, D. Z. (2018). Transient LTRE analysis reveals the demographic and trait?mediated processes that buffer population growth. Ecology letters, 21(11), 1693-1703. |
| Ctenopharyngodon idella | Animalia | Erickson, R. A., Eager, E. A., Kocovsky, P. M., Glover, D. C., Kallis, J. L., & Long, K. R. (2018). A spatially discrete, integral projection model and its application to invasive carp. Ecological Modelling, 387, 163-171. |
| NYD; 15 as Chu&Adler 2015 | Plantae | Tredennick, A., Teller, B. J., Adler, P. B., Hooker, G., & Ellner, S. P. (2018). Size-by-environment interactions: a neglected dimension of species' responses to environmental variation. bioRxiv, 329771. |
| Carduus nutans | Plantae | Hindle, B. J., Rees, M., Sheppard, A. W., Quintana?Ascencio, P. F., Menges, E. S., & Childs, D. Z. (2017). Exploring population responses to environmental change when there is never enough data: a factor analytic approach. Methods in Ecology and Evolution. |
| Eryngium cuneifolium | Plantae | Hindle, B. J., Rees, M., Sheppard, A. W., Quintana?Ascencio, P. F., Menges, E. S., & Childs, D. Z. (2017). Exploring population responses to environmental change when there is never enough data: a factor analytic approach. Methods in Ecology and Evolution. |
| Ostrea lurida | Animalia | Kimbro, D. L., White, J. W., & Grosholz, E. D. (2018). The dynamics of open populations: integration of top–down, bottom–up and supply?side influences on intertidal oysters. Oikos. |
| Gadus morua | Animalia | Färber, L., Durant, J. M., Vindenes, Y., & Langangen, Ø. (2018). Increased early offspring growth can offset the costs of long-distance spawning migration in fish. Marine Ecology Progress Series, 600, 141-150. |

| | | |
|---|---|---|
| Limosa limosa limosa | Animalia | Kentie, R., Coulson, T., Hooijmeijer, J. C., Howison, R. A., Loonstra, A. J., Verhoeven, M. A., ... & Piersma, T. (2018). Warming springs and habitat alteration interact to impact timing of breeding and population dynamics in a migratory bird. Global change biology. |
| Astragalus utahensis | Plantae | Baer, K., & Maron, J. (2018). Declining demographic performance and dispersal limitation influence the geographic distribution of the perennial forb, Astragalus utahensis (fabaceae). Journal of Ecology. |
| Helianthemum caputfelis | Plantae | Sulis, E., Bacchetta, G., Cogoni, D., & Fenu, G. (2018). Short-term population dynamics of Helianthemum caput-felis, a perennial Mediterranean coastal plant: a key element for an effective conservation programme. Systematics and Biodiversity, 1-10. |
| Aconitum noveboracense | Plantae | Louthan, A., & Doak, D. (2018). Measurement error of state variables creates substantial bias in results of demographic population models. Ecology, 99(10), 2308-2317. |
| Salvia nubicola | Plantae | Dostálek, T., Rokaya, M. B., & Münzbergová, Z. (2018). Altitude, habitat type and herbivore damage interact in their effects on plant population dynamics. PloS one, 13(12), e0209149. |
| Plantago lanceolata | Plantae | Chen, S. (2018) Modelling plant populations - do clonality and seed ecology matter? A study on Plantago lanceolata using integral projection models (IPMs). University of Sydney |
| Heracleum mantegazzianum | Plantae | Drake, J. P. (2019). An Integral Projection Model for Giant Hogweed (Master's thesis, University of Waterloo). |
| Suricata suricatta | Animalia | Paniw, M., Maag, N., Cozzi, G., Clutton-Brock, T., & Ozgul, A. (2019). Life history responses of meerkats to seasonal changes in extreme environments. Science, 363(6427), 631-635. |
| Fallopia japonica | Plantae | Lavallée, F., Smadi, C., Alvarez, I., Reineking, B., Martin, F. M., Dommanget, F., & Martin, S. (2019). A stochastic individual based model for the growth of a stand of Japanese knotweed including mowing as a management technique. arXiv preprint arXiv:1902.06971. |
| Corallium rubrum | Animalia | Montero?Serra, I., Garrabou, J., Doak, D. F., Ledoux, J. B., & Linares, C. Marine protected areas enhance structural complexity but do not buffer the consequences of ocean warming for an overexploited precious coral. Journal of Applied Ecology. |
| Ailanthus altissima | Plantae | Levin, S. C., Crandall, R. M., & Knight, T. M. (2019). Population projection models for 14 alien plant species in the presence and absence of above?ground competition. Ecology, e02681. |
| Euonymus alatus | Plantae | Levin, S. C., Crandall, R. M., & Knight, T. M. (2019). Population projection models for 14 alien plant species in the presence and absence of above?ground competition. Ecology, e02681. |
| Ligustrum obtusifolium | Plantae | Levin, S. C., Crandall, R. M., & Knight, T. M. (2019). Population projection models for 14 alien plant species in the presence and absence of above?ground competition. Ecology, e02681. |
| Lonicera maackii | Plantae | Levin, S. C., Crandall, R. M., & Knight, T. M. (2019). Population projection models for 14 alien plant species in the presence and absence of above?ground competition. Ecology, e02681. |
| Microchloa kunthii | Plantae | Martorell, C., & Martínez?Ballesté, A. (2019). Stress and disturbance determine the demographic dynamics of two grass species along a grazing gradient in Southern Mexico. Population Ecology, 61(2), 160-170. |
| Dianthus morisianus | Plantae | Cogoni, D., Sulis, E., Bacchetta, G., & Fenu, G. (2019). The unpredictable fate of the single population of a threatened narrow endemic Mediterranean plant. Biodiversity and Conservation, 1-15. |

| | | |
|---|---|---|
| Euterpe precatoria | Plantae | Isaza, C., Bernal, R., Galeano, G., & Martorell, C. (2017). Demography of Euterpe precatoria and Mauritia flexuosa in the Amazon: application of integral projection models for their harvest. Biotropica, 49(5), 653-664. |
| Mauritia flexuosa | Plantae | Isaza, C., Bernal, R., Galeano, G., & Martorell, C. (2017). Demography of Euterpe precatoria and Mauritia flexuosa in the Amazon: application of integral projection models for their harvest. Biotropica, 49(5), 653-664. |
| Erythranthe cardinalis | Plantae | Sheth, S. N., & Angert, A. L. (2018). Demographic compensation does not rescue populations at a trailing range edge. Proceedings of the National Academy of Sciences, 115(10), 2413-2418. |
| Rhizoglyphus robini | Animalia | Deere, J. A., Coulson, T., Cubaynes, S., & Smallegange, I. M. (2017). Unsuccessful dispersal affects life history characteristics of natal populations: The role of dispersal related variation in vital rates. Ecological Modelling, 366, 37-47. |
| Plesiastrea versipora | Animalia | Precoda, K., Baird, A. H., Madsen, A., Mizerek, T., Sommer, B., Su, S. N., & Madin, J. S. (2018). How does a widespread reef coral maintain a population in an isolated environment?. Marine Ecology Progress Series, 594, 85-94. |
| Hypericum cumulicola | Plantae | Quintana?Ascencio, P. F., Koontz, S. M., Smith, S. A., Sclater, V. L., David, A. S., & Menges, E. S. (2018). Predicting landscape?level distribution and abundance: Integrating demography, fire, elevation and landscape habitat configuration. Journal of Ecology, 106(6), 2395-2408. |
| Astragalus utahensis | Plantae | Baer, K. C., & Maron, J. L. (2018). Pre?dispersal seed predation and pollen limitation constrain population growth across the geographic distribution of Astragalus utahensis. Journal of Ecology, 106(4), 1646-1659. |
| Carapa guianensis | Plantae | Klimas, C. M., Cropper Jr, W. P., Kainer, K. A., & de Oliveira Wadt, L. H. (2017). Multimodel Projections for Evaluating Sustainable Timber and Seed Harvest of Carapa guianensis. Forest Science, 64(1), 15-27. |
| Arisaema triphyllum | Plantae | Heckel, C. (2015). The influence of indirect effects of large herbivores on the life history and population dynamics of unpalatable forest herb species (Doctoral dissertation, University of Pittsburgh). |
| Aurinia saxatilis ssp saxatilis | Plantae | Šimáková, T. (2018). Population biology of rock outcrop plant Aurinia saxatilis ssp saxatilis. |
| Helianthemum squamatum | Plantae | Tye, M. (2014). Integral projection models reveal interactive effects of biotic factors and disturbance on plant demography. |
| Poulsenia armata | Plantae | Zambrano, J., & Salguero?Gómez, R. (2014). Forest fragmentation alters the population dynamics of a late?successional tropical tree. Biotropica, 46(5), 556-564. |
| Fumana procumbens | Plantae | Edelfeldt, S., Bengtsson, K., & Dahlgren, J. P. (2019). Demographic senescence and effects on population dynamics of a perennial plant. Ecology, e02742. |
| Carphephorus bellidifolius | Plantae | Caughlin, T. T., Damschen, E. I., Haddad, N. M., Levey, D. J., Warneke, C., & Brudvig, L. A. (2019). Landscape heterogeneity is key to forecasting outcomes of plant reintroduction. Ecological Applications, 29(2), e01850. |
| Liatris squarrulosa | Plantae | Caughlin, T. T., Damschen, E. I., Haddad, N. M., Levey, D. J., Warneke, C., & Brudvig, L. A. (2019). Landscape heterogeneity is key to forecasting outcomes of plant reintroduction. Ecological Applications, 29(2), e01850. |
| Gaillardia aristata | Plantae | Hegstad, R. J., & Maron, J. L. (2019). Productivity and related soil properties mediate the population?level consequences of rodent seed predation on Blanketflower, Gaillardia aristata. Journal of Ecology, 107(1), 34-44. |
| Dioon caputoi | Plantae | Cabrera?Toledo, D., González?Astorga, J., Vovides, A. P., Casas, A., Vargas?Ponce, O., Carrillo?Reyes, P., ... & Vega, E. (2019). Surviving background extinction: Inferences from historic and current dynamics in the contrasting population structures of two endemic Mexican cycads. Population Ecology, 61(1), 62-73. |

| | | |
|---|---|---|
| Dioon Planifolium | Plantae | Cabrera?Toledo, D., González?Astorga, J., Vovides, A. P., Casas, A., Vargas?Ponce, O., Carrillo?Reyes, P., ... & Vega, E. (2019). Surviving background extinction: Inferences from historic and current dynamics in the contrasting population structures of two endemic Mexican cycads. Population Ecology, 61(1), 62-73. |
| Canis lupus | Animalia | Horne, J. S., Ausband, D. E., Hurley, M. A., Struthers, J., Berg, J. E., & Groth, K. (2019). Integrated population model to improve knowledge and management of Idaho wolves. The Journal of Wildlife Management, 83(1), 32-42. |
| Agelaius tricolor | Animalia | Robinson, O. J., Ruiz-Gutierrez, V., Fink, D., Meese, R. J., Holyoak, M., & Cooch, E. G. (2018). Using citizen science data in integrated population models to inform conservation. Biological Conservation, 227, 361-368. |
| Artemisia tripartita | Plantae | Adler, P. B., Kleinhesselink, A., Hooker, G., Taylor, J. B., Teller, B., & Ellner, S. P. (2018). Weak interspecific interactions in a sagebrush steppe? Conflicting evidence from observations and experiments. Ecology, 99(7), 1621-1632. |
| Hesperostipa comata | Plantae | Adler, P. B., Kleinhesselink, A., Hooker, G., Taylor, J. B., Teller, B., & Ellner, S. P. (2018). Weak interspecific interactions in a sagebrush steppe? Conflicting evidence from observations and experiments. Ecology, 99(7), 1621-1632. |
| Pseudoroegneria spicata | Plantae | Adler, P. B., Kleinhesselink, A., Hooker, G., Taylor, J. B., Teller, B., & Ellner, S. P. (2018). Weak interspecific interactions in a sagebrush steppe? Conflicting evidence from observations and experiments. Ecology, 99(7), 1621-1632. |
| Poa secunda | Plantae | Adler, P. B., Kleinhesselink, A., Hooker, G., Taylor, J. B., Teller, B., & Ellner, S. P. (2018). Weak interspecific interactions in a sagebrush steppe? Conflicting evidence from observations and experiments. Ecology, 99(7), 1621-1632. |
| Rhizoglyphus robini | Animalia | Smallegange, I. M., & Ens, H. M. (2018). Trait?based predictions and responses from laboratory mite populations to harvesting in stochastic environments. Journal of Animal Ecology, 87(4), 893-905. |
| Chamaedorea elegans | Plantae | Jansen, M., Anten, N. P., Bongers, F., Martínez?Ramos, M., & Zuidema, P. A. (2018). Towards smarter harvesting from natural palm populations by sparing the individuals that contribute most to population growth or productivity. Journal of applied ecology, 55(4), 1682-1691. |
| Tridacna maxima | Animalia | Schreiber, S. J., & Moore, J. L. (2018). The structured demography of open populations in fluctuating environments. Methods in Ecology and Evolution, 9(6), 1569-1580. |
| Psidium cattleianum | Plantae | Horvitz, C. C., Denslow, J. S., Johnson, T., Gaoue, O., & Uowolo, A. (2018). Unexplained variability among spatial replicates in transient elasticity: implications for evolutionary ecology and management of invasive species. Population ecology, 60(1-2), 61-75. |
| Asclepias curassavica | Plantae | Kellett, K. M., & Shefferson, R. P. (2018). Temporal variation in reproductive costs and payoffs shapes the flowering strategy of a neotropical milkweed, Asclepias curassavica. Population ecology, 60(1-2), 77-87. |
| Ailanthus altissima | Plantae | Crandall, R. M., & Knight, T. M. (2018). Role of multiple invasion mechanisms and their interaction in regulating the population dynamics of an exotic tree. Journal of applied ecology, 55(2), 885-894. |
| Prioria copaifera | Plantae | Needham, J., Merow, C., Chang-Yang, C. H., Caswell, H., & McMahon, S. M. (2018). Inferring forest fate from demographic data: from vital rates to population dynamic models. Proceedings of the Royal Society B: Biological Sciences, 285(1874), 20172050. |
| Calophyllum longifolium | Plantae | Needham, J., Merow, C., Chang-Yang, C. H., Caswell, H., & McMahon, S. M. (2018). Inferring forest fate from demographic data: from vital rates to population dynamic models. Proceedings of the Royal Society B: Biological Sciences, 285(1874), 20172050. |

| | | |
|---|---|---|
| Garcinia intermedia | Plantae | Needham, J., Merow, C., Chang-Yang, C. H., Caswell, H., & McMahon, S. M. (2018). Inferring forest fate from demographic data: from vital rates to population dynamic models. Proceedings of the Royal Society B: Biological Sciences, 285(1874), 20172050. |
| Upupa epops | Plantae | Plard, F., Schindler, S., Arlettaz, R., & Schaub, M. (2018). Sex-specific heterogeneity in fixed morphological traits influences individual fitness in a monogamous bird population. The American Naturalist, 191(1), 106-119. |
| Ursus arctos | Animalia | Bled, F., Belant, J. L., Van Daele, L. J., Svoboda, N., Gustine, D., Hilderbrand, G., & Barnes Jr, V. G. (2017). Using multiple data types and integrated population models to improve our knowledge of apex predator population dynamics. Ecology and evolution, 7(22), 9531-9543. |
| 33 tree species | Plantae | Visser, M. D., Schnitzer, S. A., Muller?Landau, H. C., Jongejans, E., de Kroon, H., Comita, L. S., ... & Wright, S. J. (2018). Tree species vary widely in their tolerance for liana infestation: A case study of differential host response to generalist parasites. journal of ecology, 106(2), 781-794. |
| Bertholletia Excelsa | Plantae | Bertwell, T. D., Kainer, K. A., Cropper Jr, W. P., Staudhammer, C. L., & de Oliveira Wadt, L. H. (2018). Are Brazil nut populations threatened by fruit harvest?. Biotropica, 50(1), 50-59. |
| Dendroctonus ponderosae | Animalia | Goodsman, D. W., Aukema, B. H., McDowell, N. G., Middleton, R. S., & Xu, C. (2018). Incorporating variability in simulations of seasonally forced phenology using integral projection models. Ecology and evolution, 8(1), 162-175. |
| Schiedea obovata | Plantae | Bialic?Murphy, L., & Gaoue, O. G. (2018). Low interannual precipitation has a greater negative effect than seedling herbivory on the population dynamics of a short?lived shrub, Schiedea obovata. Ecology and evolution, 8(1), 176-184. |
| Hibiscus meyeri | Plantae | Louthan, A. M., Pringle, R. M., Goheen, J. R., Palmer, T. M., Morris, W. F., & Doak, D. F. (2018). Aridity weakens population-level effects of multiple species interactions on Hibiscus meyeri. Proceedings of the National Academy of Sciences, 115(3), 543-548. |
| Cirsium vulgare | Plantae | Schultz, E. L. (2018) Populations in a changing world: the effects of environmental variation on species with complex life histories. Rice University |
| Hypericum cumulicola | Plantae | Quintana?Ascencio, P. F., Koontz, S. M., Ochocki, B., Sclater, V. L., López?Borghesi, F., Li, H., & Menges, E. S. Assessing the roles of seed bank, seed dispersal and historical disturbances for metapopulation persistence of a pyrogenic herb. Journal of Ecology. |
| Veratrum album | Plantae | Franco, D., Guiver, C., Logemann, H., & Perán, J. (2019). Boundedness, persistence and stability for classes of forced difference equations arising in population ecology. Journal of Mathematical Biology, 1-48. |
| Ipomopsis aggregata | Plantae | Campbell, D. R. (2019). Early snowmelt projected to cause population decline in a subalpine plant. Proceedings of the National Academy of Sciences, 201820096. |
| Ipomopsis tenuituba | Plantae | Campbell, D. R. (2019). Early snowmelt projected to cause population decline in a subalpine plant. Proceedings of the National Academy of Sciences, 201820096. |
| Boswellia papyrifera | Plantae | Bongers, F., Groenendijk, P., Bekele, T., Birhane, E., Damtew, A., Decuyper, M., ... & Lemenih, M. (2019). Frankincense in peril. Nature Sustainability, 2, 602-610. |
| Sebastes mystinus | Animalia | Nickols, K. J., White, J. W., Malone, D., Carr, M. H., Starr, R. M., Baskett, M. L., ... & Botsford, L. W. (2019). Setting ecological expectations for adaptive management of marine protected areas. Journal of Applied Ecology. |
| Thamnophis gigas | Animalia | |
| Yermo xanthocephalus | Plantae | Dibner, R.R, Peterson, M. L., Louthan, A. M., & Doak, D.F. (2019). Multiple mechanisms confer stability to isolated populations of a rare endemic plant. Ecological Monographs 89 (2): e01360. |

| | | |
|---|---|---|
| Bugula neritina | Animalia | Cameron, H., Coulson, T., & Marshall, D. J. (2019) Size and density mediate transitions between competition and facilitation. Ecology Letters. DOI: 10.1111/ele.13381 |
| Watersipora subtorquata | Animalia | Cameron, H., Coulson, T., & Marshall, D. J. (2019) Size and density mediate transitions between competition and facilitation. Ecology Letters. DOI: 10.1111/ele.13381 |
| Arisaema triphyllum | Plantae | Bialic-Murphy, L., Heckel, C.D., McElderry, R.M., & Kalisz, S. (2019) Deer indirectly alter the reproductive strategy and operational sex ratio of an unpalatable forest perennial. American Naturalist 195 (1), 56-69 |
| Poecilia reticulata | Animalia | De Bona, S. (2019) Dispersal, habitat use, and the invasion dynamics of introduced populations: A case study on the Guppy. (Doctoral Dissertation, University of Jyvaskyla) |
| Koeleria macrantha | Plantae | Collins, C.G., Bohner, T.F., & Diez J.M. (2019) Plant-soil feedbacks and faciliation influence the demography of herbaceous alpine species in response to woody plant range expansion. Frontiers in Ecology and Evolution. DOI: 10.3389/fevo.2019.00417 |
| Eriogonum ovalifolium | Plantae | Collins, C.G., Bohner, T.F., & Diez J.M. (2019) Plant-soil feedbacks and faciliation influence the demography of herbaceous alpine species in response to woody plant range expansion. Frontiers in Ecology and Evolution. DOI: 10.3389/fevo.2019.00417 |
| Esox lucius | Animalia | Stubberud, M. W., Vindenes, Y., Vollestad, L.A., Winfield, I.J., Stenseth, N. C., & Langangen, O. (2019) Effects of size- and sex-selective harvesting: An integral projection model approach. Ecology & Evolution. DOI: 10.1002/ece3.5719 |
| Rheum nobile | Plantae | Song, B., Stoll, P., Peng, D., Sun, H., & Stoecklin, J. (2019) Demography of the giant monocarpic herb Rheum nobile in the Himalayas and the effect of disturbances by grazing. Annals of Botany. DOI: 10.1093/aob/mcz178 |
| 4 coral species | Animalia | Noriega, M.A. (2019) Competition and coexistance of reef-corals (PhD Dissertation, James Cook University) |
| Ostrea edulis | Animalia | Lown, A.E. (2019) Community ecology and population dynamics of the European native oyster (Ostrea edulis) in Essex, UK: A baseline for the management of the Blackwater, Crouch, Roach and Colne Estuaries Marine Conservation Zone. (PhD Dissertation, University of Essex) |
| Pimephales promelas | Animalia | Pollesch, N.L., Flynn, K.M., Kadlec, S.M., Swintek, J.A., & Etterson, M.A. Developing integral projection models for ecotoxicology. URL: https://scse.d.umn.edu/sites/scse.d.umn.edu/files/polleschnate__paper__2__12062019.[illegible] |
| Cervus canadensis | Animalia | Lachish, S., Brandell, E., Craft, M., Dobson, A., Hudson, P., Macnulty, D., & Coulson, T. (2019) Investigating the dynamics of elk population size and body mass in a seasonal environment using a mechanistic integral projection model. Am Nat |
| Hymanaea courbaril | Plantae | Marques, I., Vidal, E., Tomazello-Filho, M., & Groenendijk, P. (2019) Applying tree rings and population models to improve future timber yield projections of Hymenaea courbaril (Jatoba) in the Eastern Amazon. Pesq. Flor. Bras. 39. |
| Daphnia magna | Animalia | Bruijning, M. (2019) Persisting in an ever-changing world: Integrating plastic and genetic responses across the life cycle. (PhD Dissertation, Radboud University Nijmegen) |
| Lathyrus vernus | Plantae | Greiser, C., Hylander, K., Meineri, E., Luoto, M., & Ehrlen, J. (2020) Climate limitation at the cold edge: contrasting perspectives from species distribution modelling and a transplant experiment. Ecography 43: 1-11. |
| Echinomastus erectocentrus var. acunensis | Plantae | Larios, E., Gonzalez, E.J., Rosen, P.C., Pate, A., & Holm, P. (2020) Population projections of an endangered cactus suggest little impact of climate change. Oecologia. DOI: 10.1007/s00442-020-04595-y |
| Rhizoglyphus robini | Animalia | Deere, J.A., van den Berg, I., Roth, G., Smallegange, I.M. (2020) Modelling the impact of dispersal on a natal population that exhibits boom-bust dynamics. bioRxiv. DOI: 10.1101/402198 |

| | | |
|---|---|---|
| Ovis aries | Animalia | Kentie, R., Clegg, S.M., Tuljapurkar, S., Gaillard, J-M., Coulson, T. (2020) Life-history strategy varies with the strength of competition in a food-limited ungulate population. Ecology Letters. DOI: 10.1111/ele.13470 |
| Actea spicata | Plantae | Roemer, G., Christiansen, D.M., de Buhr, H., Hylander, K., Jones, O.R., Merinero, S., Reitzel, K., Ehrlen, J., & Dahlgren, J.P. (2020). Drivers of large-scale spatial demographic variation in a perennial plant. DOI: 10.1101/2020.02.29.969428 |
| Podarcis lilfordi | Animalia | Rotger, A., Igual, J.M., & Tavecchia, G. (2020) Contrastic size-dependent life history strategies of an insular lizard. Current Zoology 2020: 1-9. DOI: 10.1093/cz/zoaa019 |
| Ambystoma bishopi | Animalia | Brooks, G.M. (2020) Chapter 4: Population viability of an endangered amphibian under future management scenarios, In: On the use of demographic models to inform amphibian conservation and management: a case study of the Reticulated Flatwoods Salamander. |
| Branta leucopsis | Animalia | Layton-Matthews, K. (2020) Demographic consequences of rapid climate change and density dependence in migratory Arctic Geese. PhD Thesis. |
| Nerodia sipedon | Animalia | Rose, J.P., & Todd, B.D. (2020) Targeting eradication of introduced watersnakes using integral projection models. Animal Conservation. DOI: 10.1111/acv.12590 |
| Pseudoroegneria spicata | Plantae | Shriver, R.K., Campbell, E., Dailey, C., Gaya, H., Hill, A., Kuzminski, S., Miller-Bartley, M., Moen, K., Moettus, R., Oschrin, E., Reese, D., Simonson, M., Willson, A, & Parker, T.H. (2020) Local landscape position impacts demographic rates in a widespread North American steppe bunchgrass. DOI: 10.32942/osf.io/8pdwg |
| Lupinus polyphyllus | Plantae | Ramula S. (2020) Annual mowing has the potential to reduce the invasion of herbaceous Lupinus polyphyllus. Biological lnvasions 22: 3163-3173. |
| Cyclindriopuntia imbricata | Plantae | Czachura, K., & Miller T.E.X. (2020) Demographic back-casting reveals that subtle dimensions of climate change have strong effects on population viability. Journal of Ecology. DOI: 10.1111/1365-2745.13471 |
| Lupinus tidestromii | Plantae | Compagnoni A., Pardini E., & Knight T.M. (2020) Increasing temperature threatens an already endangered coastal dune plant. bioRxiv. DOI: 10.1101/2020.08.02.2333288 |
| Porites divaricata | Animalia | Lord K.S. (2020) The importance of mangroves as habitat for corals. Reef Encounter 35(1): 36-40. |
| Sebastes auriculatus | Animalia | Perkins, N.R., Prall, M., Chakraborty A., White, J.W., Baskett, M.L., & Morgan S.G. (2020) Quantifying the statistical power of monitoring programs for marine protected areas. Ecological Applications. DOI: 10.1002/eap.2215 |
| Ranunculus austro-oreganus | Plantae | Reed, P.B., Peterson, M.L., Pfeifer-Meister, L.E., Morris, W.F., Doak, D.F., Roy, B.A., Johnson, B.R., Bailes, G.T., Nelson, A.A., & Bridgham, S.D. (2020) Climate manipulations differentially affect plant population dynamics within versus beyond northern range limits. Journal of Ecology. DOI: 10.1111/1365-2745.13494 |
| Sidelcea malviflora | Plantae | Reed, P.B., Peterson, M.L., Pfeifer-Meister, L.E., Morris, W.F., Doak, D.F., Roy, B.A., Johnson, B.R., Bailes, G.T., Nelson, A.A., & Bridgham, S.D. (2020) Climate manipulations differentially affect plant population dynamics within versus beyond northern range limits. Journal of Ecology. DOI: 10.1111/1365-2745.13494 |
| Microseris laciniata | Plantae | Reed, P.B., Peterson, M.L., Pfeifer-Meister, L.E., Morris, W.F., Doak, D.F., Roy, B.A., Johnson, B.R., Bailes, G.T., Nelson, A.A., & Bridgham, S.D. (2020) Climate manipulations differentially affect plant population dynamics within versus beyond northern range limits. Journal of Ecology. DOI: 10.1111/1365-2745.13494 |
| Achnatherum lemmonii | Plantae | Reed, P.B., Peterson, M.L., Pfeifer-Meister, L.E., Morris, W.F., Doak, D.F., Roy, B.A., Johnson, B.R., Bailes, G.T., Nelson, A.A., & Bridgham, S.D. (2020) Climate manipulations differentially affect plant population dynamics within versus beyond northern range limits. Journal of Ecology. DOI: 10.1111/1365-2745.13494 |

| | | |
|---|---|---|
| Festuca roemeri | Plantae | Reed, P.B., Peterson, M.L., Pfeifer-Meister, L.E., Morris, W.F., Doak, D.F., Roy, B.A., Johnson, B.R., Bailes, G.T., Nelson, A.A., & Bridgham, S.D. (2020) Climate manipulations differentially affect plant population dynamics within versus beyond northern range limits. Journal of Ecology. DOI: 10.1111/1365-2745.13494 |
| Danthonia californica | Plantae | Reed, P.B., Peterson, M.L., Pfeifer-Meister, L.E., Morris, W.F., Doak, D.F., Roy, B.A., Johnson, B.R., Bailes, G.T., Nelson, A.A., & Bridgham, S.D. (2020) Climate manipulations differentially affect plant population dynamics within versus beyond northern range limits. Journal of Ecology. DOI: 10.1111/1365-2745.13494 |
